# A dual histone code specifies the binding of heterochromatin protein Rhino to a subset of piRNA source loci

**DOI:** 10.1101/2024.01.11.575256

**Authors:** Abdou Akkouche, Emma Kneuss, Susanne Bornelöv, Yoan Renaud, Evelyn L. Eastwood, Jasper van Lopik, Nathalie Gueguen, Mingxuan Jiang, Pau Creixell, Stéphanie Maupetit-Mehouas, Anna Sobieszek, Yifan Gui, Benjamin Czech Nicholson, Emilie Brasset, Gregory J. Hannon

**Author notes:** To whom correspondence should be addressed: BCN, EB, GJH. These authors contributed equally (Abdou Akkouche, Emma Kneuss, Susanne Bornelöv).

## Abstract

Animal germ cells deploy a specialized small RNA-based silencing system, called the PIWI-interacting RNA (piRNA) pathway, to prevent unwanted expression of transposable elements and maintain genome integrity. In *Drosophila* germ cells, the majority of piRNA populations originate from dual-strand piRNA clusters, genomic regions highly enriched in transposable element (TE) fragments, via an elaborate protein machinery centred on the heterochromatin protein 1 homolog, Rhino. Although Rhino binds to peptides carrying trimethylated H3K9 *in vitro*, it is not fully understood why *in vivo* only a fraction of H3K9me3-decorated heterochromatin is occupied by Rhino. Recent work revealed that Rhino is recruited to a subset of piRNA clusters by the zinc finger protein Kipferl. Here we identify a Kipferl-independent mode of Rhino targeting that, in addition to the previously established role of H3K9me3, also depends on the histone H3 lysine 27 methyltransferase Enhancer of Zeste. At Kipferl-independent sites, we find that Rhino, through its chromodomain, specifically binds to loci marked by both H3K9me3 and H3K27me3. Although the exact mechanism of how Rhino binding is influenced by dual histone modifications remains unclear from a structural and biochemical perspective, our work suggests that combinatorial modifications can play a crucial role in influencing the specificity of chromatin-binding protein interactions. These findings provide an enhanced understanding of the multifaceted mechanisms by which Rhino targets piRNA source loci highlighting the sophisticated epigenetic landscape governing TE silencing in *Drosophila* germ cells. Our work further reveals a role for dual histone modifications defining the binding specificity of a key chromatin protein.

## INTRODUCTION

Transposable elements (TEs) are genetic sequences, present in nearly all organisms, that have the ability to move and insert themselves into different positions within a genome. While TEs can contribute to genetic diversity and evolution, their uncontrolled activity in gonadal cells can have detrimental effects, including the potential to dramatically reduce fertility. Thus, in animal gonads, a dedicated mechanism known as the PIWI-interacting RNA (piRNA) pathway plays a crucial role in silencing and controlling active TEs to safeguard germline integrity (Brennecke et al. 2007; Czech et al. 2018; Ozata et al. 2019). The piRNA machinery utilises 23-to 30-nt small RNA molecules, which associate with PIWI family proteins to recognize and target TE transcripts for post-transcriptional degradation or co-transcriptional silencing. While some piRNAs are derived from active TEs inserted across the genome, the majority originate from dedicated piRNA source loci. Of those, a subset, called piRNA clusters, are enriched in TE content. All reported piRNA clusters in somatic follicle cells resemble canonical transcription units, give rise to piRNAs from one genomic strand, and are referred to as unistrand clusters (van Lopik et al. 2023). In contrast, piRNA clusters in the germline are typically expressed non-canonically, producing piRNAs from both genomic strands and are referred to as dual-strand clusters (Klattenhoff et al. 2009; Goriaux et al. 2014; Mohn et al. 2014; Zhang et al. 2014; Chen et al. 2016; Andersen et al. 2017).

Rhino (Rhi), a homologue of Heterochromatin Protein 1a (HP1a), is a germline-specific piRNA pathway factor essential for piRNA production, TE repression, and fertility (Klattenhoff et al. 2009). Both Rhi and HP1a belong to a distinct group of chromatin-binding proteins containing chromodomains (CDs). Among CD-containing proteins, different unique binding properties exist. For instance, *Drosophila* HP1a is primarily associated with H3K9me2/3-enriched regions such as constitutive heterochromatin, whereas the developmental regulator Polycomb (Pc), another chromodomain protein, specifically binds to H3K27me3 marks often found in facultative heterochromatin (Franke et al. 1992; Fischle et al. 2003; Bernstein et al. 2006). In line with Rhi being evolutionary related to HP1a, several studies have shown that *in vivo* Rhi specifically localises to a subset of piRNA source loci, including dual-strand piRNA clusters, which are typically characterised by the presence of H3K9me3 marks (Klattenhoff et al. 2009; Le Thomas et al. 2014; Mohn et al. 2014; Zhang et al. 2014). *In vitro*, the Rhi CD was shown to recognize di-/tri-methylated H3K9 peptides (Le Thomas et al. 2014; Mohn et al. 2014; Yu and Huang 2015). Disruption of the H3K9 methyltransferase Eggless (Egg) compromises piRNA cluster transcription, highlighting a role of H3K9me3 in Rhi recruitment to piRNA clusters (Rangan et al., 2011).

At the genomic regions to which it binds, Rhi acts as nucleator for a multi-protein complex that triggers non-canonical transcription from heterochromatic regions and facilitates the export of the resulting piRNA precursors from the nucleus (Klattenhoff et al. 2009; Pane et al. 2011; Le Thomas et al. 2014; Mohn et al. 2014; Zhang et al. 2014; Chen et al. 2016; Hur et al. 2016; Andersen et al. 2017; Zhang et al. 2018; ElMaghraby et al. 2019; Kneuss et al. 2019; Zhang et al. 2021). Although it was demonstrated that Rhi binds the repressive histone mark H3K9me3 *in vitro* (Le Thomas et al. 2014; Mohn et al. 2014; Yu and Huang 2015), Rhi immunofluorescence staining and previously published ChIP-seq data indicate that *in vivo* Rhi associates only with a fraction of genomic regions enriched in H3K9me3 marks (Mohn et al. 2014), hence raising questions about how its specificity is determined *in vivo*. A recent study identified the zinc finger domain protein Kipferl (Kipf) as an essential cofactor for Rhi in the female germline, with their interaction being mediated through Rhi’s CD (Baumgartner et al. 2022; Baumgartner et al. 2024). Kipf was shown to recruit Rhi to dedicated heterochromatic loci that exhibit enrichment in a guanosine-rich motif, suggesting that sequence content within these regions could contribute to the binding specificity of Rhi. However, while the interaction between Kipf and Rhi is critical for the recruitment of Rhi to numerous genomic loci, prominent Rhi-dependent piRNA producing regions, such as piRNA clusters *42AB* and *38C*, show little to no dependence on Kipf (Baumgartner et al. 2022). These observations suggest that additional cues mediate and/or contribute to Rhi’s chromatin binding pattern. Here we uncover an unexpected, Kipf-independent mechanism for Rhi targeting that relies on the H3K27me3 methyltransferase E(z). We propose a model in which dual decoding of H3K9me3 and H3K27me3 guides Rhino to a subset of piRNA source loci, including the most prominent cluster *42AB*.

## RESULTS

### The histone methyltransferase E(z) is required for TE silencing in *Drosophila* germ cells

Understanding the mechanisms by which Rhi is recruited to and interacts with chromatin at piRNA source loci is crucial for unravelling the complexities of TE silencing and genome integrity. We therefore searched for candidate factors that could affect TE control in germ cells through a focused reverse-genetic screen in *Drosophila* ovaries (**Fig. 1a**). We specifically targeted chromatin proteins as well as histone modifiers and probed the effects of their depletion on the expression of germline-specific TEs by quantitative RT-PCR (RT-qPCR). Knockdown of 32 of these genes in germ cells resulted in TE de-repression and highlighted the potential role of different protein complexes in TE regulation (e.g., COMPASS complex, NSL complex) (**Extended Data Fig. 1**).

**Figure 1:**
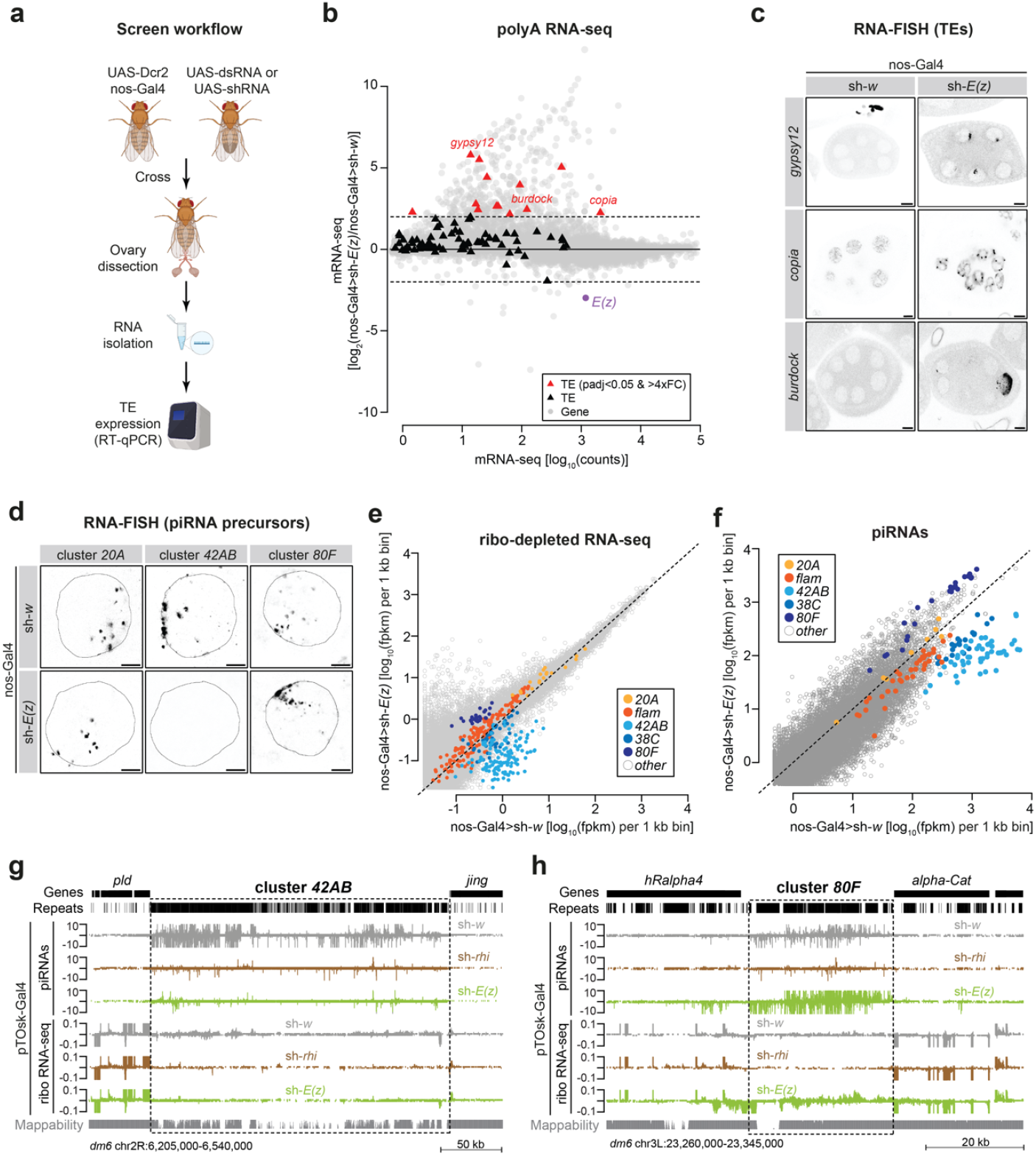
E(z) is required for TE silencing in *Drosophila* germ cells and affects piRNA production. **a**, Schematic showing the *in vivo* screen workflow. Male flies carrying dsRNA or shRNA constructs under the inducible UAS promoter were crossed to females expressing the germline-specific nos-GAL4 driver and a *Dcr2* transgene. F1 offspring were transferred to new vials and fertility was determined by counting eggs laid and hatched. RNA was extracted from ovaries of F1 offspring and TE levels were measured by RT-qPCR. **b**, MA plot showing counts per gene (grey) and TEs (black) in polyA-selected RNA-seq libraries from ovaries depleted of w (control) and E(z) using the nos-Gal4 driver. TEs with padj<0.05 and >4x fold-change are shown in red, E(z) is indicated in purple. **c**, Confocal images showing RNA-FISH signal for the indicated TEs (*burdock, copia*, and *gypsy12*) upon nos-Gal4 mediated knockdown or *w* or *E(z)* (scale bar: 10 µm). **d**, Confocal images showing RNA-FISH signal for transcripts derived from piRNA clusters *42AB, 80F* and *20A* in control and *E(z)* depleted ovaries using the nos-Gal4 driver (scale bar: 5 μm). **e**, Scatter plot depicting normalized ribo-depleted RNA levels (fpkm) of uniquely mapping reads in 1 kb bins in ovaries with nos-Gal4 driven *E(z)* knockdown versus control (average of three replicate experiments each). **f**, Scatter plot depicting normalized piRNA levels of uniquely mapping piRNAs (fpkm) in 1 kb bins in ovaries with nos-Gal4 driven *E(z)* knockdown versus control (two replicates each). **g**, UCSC genome browser tracks displaying the dual-strand piRNA cluster *42AB*. Levels of uniquely mapping piRNAs and ribo-depleted RNAs from ovaries with the indicated knockdowns using the pTOsk-Gal4 driver (average of three replicate experiments each) are shown along with tracks displaying genes, repeats and mappability. Dashed line indicates approximate piRNA cluster boundaries. **h**, as in g but showing dual-strand piRNA cluster *80F*

As expected from previous reports (Rangan et al. 2011), depletion of the H3K9me3 methyltransferase Egg resulted in a strong deregulation of germline TEs and was accompanied by loss of Rhi foci by immunofluorescence (**Extended Data Fig. 1, Extended Data Fig. 2a)**. Unexpectedly, however, germline-specific depletion of the histone methyltransferase Enhancer of Zeste (E(z)), resulted in upregulated TE levels (**Extended Data Fig. 1**). Of note, we also found that the depletion of other subunits of the Polycomb repressive complex 2 (PRC2), namely *su(z)12* and *caf1-55*, resulted in the de-repression of some germline TEs (see **Supplementary Note 1**). In *Drosophila*, E(z) is responsible for trimethylation of lysine 27 on histone H3 (H3K27me3) and is essential for Polycomb-mediated chromatin silencing (Margueron and Reinberg 2011; DeLuca et al. 2020). We found that depletion of E(z) using a strong Gal4 driver (UAS-Dcr2 transgene combined with nos-Gal4, referred to as D2G4) caused rudimentary ovaries. To exclude the possibility that the observed TE de-repression was due to pleiotropic oogenesis defects, we used different Gal4 drivers, namely nos-Gal4 which is active from the germarium through stage 2, then inactive between stages 3 and 6 of oogenesis but reactivated at later stages (Van Doren et al. 1998), and TOsk-Gal4 which is expressed immediately after germline cyst formation (ElMaghraby et al. 2022), as well as two independent E(z)-RNAi lines, all of which resulted in nearly wildtype ovarian morphology despite reduced H3K27me3 levels in nurse cell nuclei (**Extended Data Fig. 2b**), and unchanged Rhi localisation as observed by immunofluorescence (**Extended Data Fig. 2a**). Sequencing of polyA-containing RNAs from ovaries depleted of *E(z)* using the nos-Gal4 driver revealed a strong upregulation of 13 germline TE families (fold change [FC]>4, padj<0.05), including *gypsy12, burdock* and *copia* (**Fig. 1b)**. TE de-repression results were validated by RNA-FISH, with *burdock, gypsy12* and *copia* RNA levels strongly elevated in germ cells of ovaries depleted of *E(z)* (**Fig. 1c**). Overall, however, the effects on TEs that we observed for *E(z)* knockdown were less severe compared to ovaries depleted of the piRNA factor Rhi, which affected 53 TE families (FC>4, padj<0.05), suggestive of partial contributions of E(z) to Rhi function (**Extended Data Fig. 2c**; data from (Baumgartner et al. 2022)). To distinguish between compromised global chromatin silencing and a specific role in the piRNA pathway, we performed RNA-FISH for precursor transcripts derived from the dual-strand piRNA clusters *42AB* (Kipf-independent) and *80F* (Kipf-dependent), and the unistrand cluster *20A* (germline-enriched but Rhi-independent). Germline-specific knockdown of *E(z)* revealed a strong loss in piRNA precursors derived from *42AB*, but not for transcripts originating from *80F* and *20A*, suggesting a potential role of E(z) in the transcription of Kipf-independent dual-strand piRNA clusters (**Fig. 1d, Extended Data Fig. 2d**).

**Figure 2:**
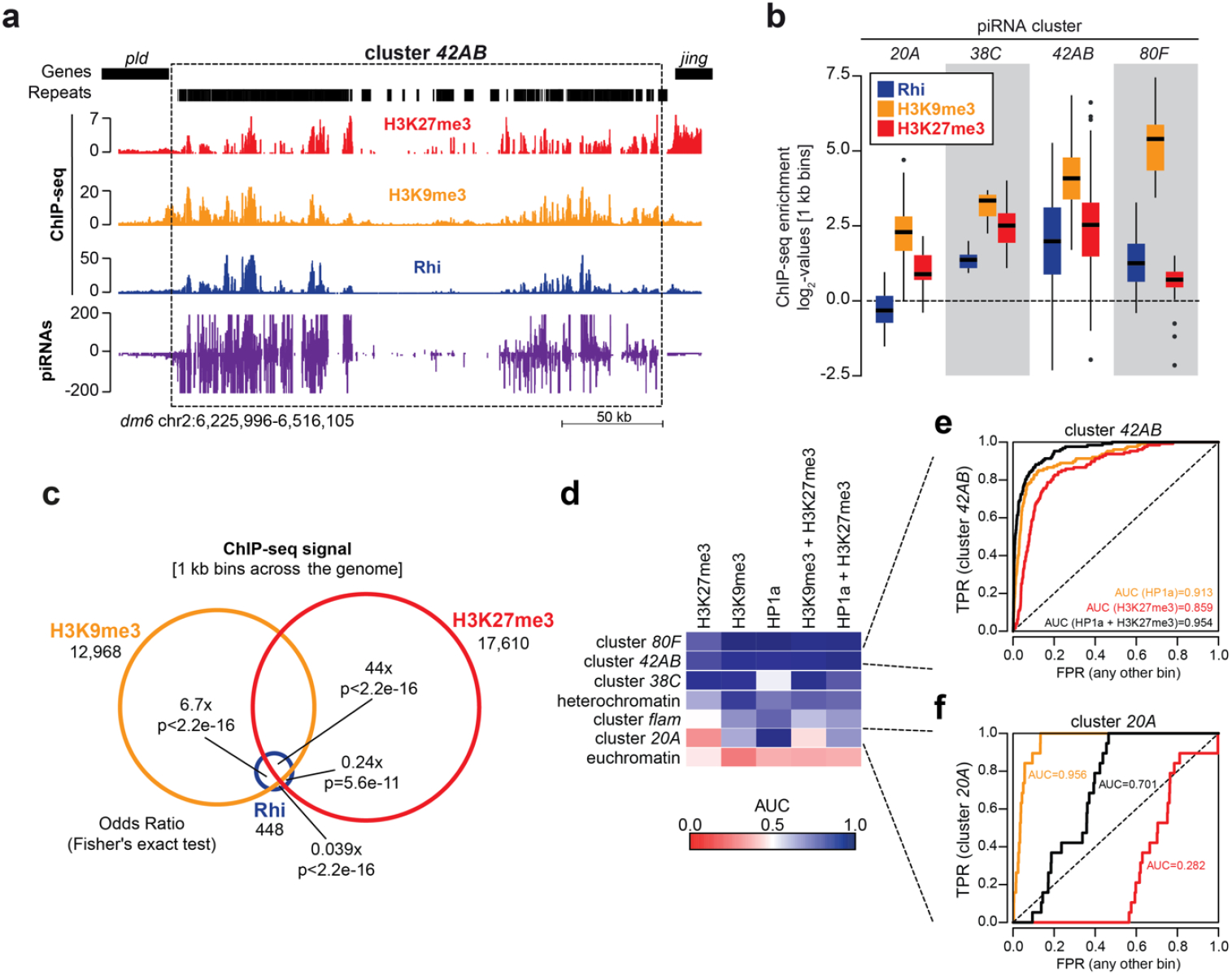
H3K27me3 is present on certain piRNA source loci. **a**, UCSC genome browser tracks of the region comprising dual-strand piRNA cluster *42AB* displaying ChIP-seq signal (depicted as coverage per million reads) of Rhi (blue), H3K27me3 (red), H3K9me3 (yellow), uniquely mapping piRNAs (purple, coverage normalized to miRNA reads), and tracks indicating genes and repeats in nos-Gal4>sh-*w* ovaries. **b**, Box plot showing average log_2_-fold ChIP-seq enrichment over input of H3K27me3 (red), H3K9me3 (yellow), and Rhi (blue) at the indicated piRNA source loci (n = 1 kb bins) in control ovaries. Box plots show median (centre line), with interquartile range (box) and whiskers indicate at most 1.5x interquartile range). **c**, Venn diagram showing overlap between Rhi, H3K9me3, and H3K27me3 ChIP-seq signal across genomic 1 kb bins. Signal was considered to be present if the bin overlapped a MACS2 broad peak (q<0.05, broad-cutoff<0.1) across two pooled biological replicates. Enrichment was calculated as an Odds Ratio (OR) between Rhi binding in the presence of indicated factor or without. **d**, Heatmap displaying predictive performance measured as area under ROC curve (AUC) across different chromatin marks (top) and target regions and (left). The regions are ordered according to their AUC for HP1a + H3K27me3. **e**, Line graph comparing the ability to identify dual-strand piRNA cluster *42AB* based on the strength of individual features (H3K27me3 or HP1a) and combinations (mean signal for H3K27me3 + HP1a). **f**, as in e but showing unistrand piRNA cluster *20A*.

Next, we analysed the effects of *E(z)* knockdown on piRNA precursors globally using ribo-depleted RNA-seq. To measure transcription across all piRNA source loci, we divided the genome into 1 kb bins and quantified the signal per bin, similar to previous work (Mohn et al. 2014). Upon germline-specific *E(z)* depletion, we observed a strong reduction in precursor transcript levels across the dual-strand cluster *42AB*, and moderately reduced levels for cluster *38C*, whereas the unistrand piRNA clusters *20A* or *flam* (soma-expressed and Rhi-independent) were essentially unchanged (**Fig. 1e, Extended Data Fig. 2e**). Strikingly, precursor levels for Kipf-dependent piRNA cluster *80F* were increased upon *E(z)* knockdown. RT-qPCR analysis confirmed the results obtained by RNA-seq (**Extended Data Fig. 2f**). Of note, and as reported previously (Klattenhoff et al. 2009; Le Thomas et al. 2014; Mohn et al. 2014), Rhi-depleted ovaries showed a specific and global reduction of precursor transcripts that originate from dual-strand clusters (**Extended Data Fig. 2g**). These data suggest a more nuanced role of E(z) in TE silencing in the germline as compared to Rhi.

Next, we sequenced piRNA populations from *Drosophila* ovaries in which E(z) or Rhi were depleted specifically in germ cells and compared these to control knockdowns (targeting the *white* gene) (**Table S1**). In agreement with the reduction of precursor transcripts from dual-strand clusters *38C* and *42AB* upon *E(z)* knockdown (**Fig. 1e**), we detected a severe reduction in mature piRNA levels produced from those loci (**Fig. 1f**). As expected from their precursor expression, piRNA levels derived from the cluster *80F* were more abundant, while piRNAs from the unistrand clusters *20A* and *flam*, were unaffected upon *E(z)* germline knockdown (**Fig. 1f, Extended Data Fig. 2h**). The reduction in precursor and piRNA levels observed at cluster *42AB* was similar for ovaries depleted for E(z) and Rhi (**Fig. 1g**). In contrast, transcript levels and piRNA expression were enhanced at the *80F* cluster upon *E(z)* knockdown, whereas there was a pronounced reduction of these RNA populations in the *rhi* knockdown (**Fig. 1h**).

Ovaries depleted of E(z) also showed a reduction in antisense piRNAs targeting individual germline-specific Ovaries depleted of E(z) also showed a reduction in antisense piRNAs targeting individual germline-specific TEs, which could explain the observed increase in their mRNA levels (**Extended Data Fig. 2i**). As expected, piRNAs against TEs that are mainly active in the somatic compartment of the ovary (e.g., *Tabor, gypsy, ZAM*), were not affected upon E(z) depletion, likely due to their origin from the soma-specific *flam* cluster (**Extended Data Fig. 2i**,**j**). Altogether, our results reveal that the histone methyltransferase E(z) is required for TE silencing and its depletion affects piRNA production in germ cells from a subset of dual-strand clusters.

### H3K27me3 and H3K9me3 marks co-exist on E(z)-dependent piRNA source loci

To elucidate why certain dual-strand piRNA clusters are dependent on E(z), we examined the genome-wide distribution of H3K27me3, which is deposited by E(z), and H3K9me3, which has been associated with piRNA clusters and Rhi recruitment, by performing ChIP-seq and CUT&RUN in *Drosophila* ovaries (**Extended Data Fig. 3a**). These experiments showed that a number of genomic regions, including the dual-strand piRNA cluster *42AB*, in addition to being decorated by H3K9me3, are also decorated with H3K27me3 (**Fig. 2a**,**b**). A similar, albeit less strong, pattern of H3K9me3 and H3K27me3 was observed at other piRNA source loci, including the *ey*/*Sox102F* region on chromosome 4 (**Extended Data Fig. 3b**). Notably, we found that Rhi preferentially occupied regions decorated both by H3K9me3 and H3K27me3 over those only carrying H3K9me3 (44-fold enrichment vs 6.7-fold enrichment) (**Fig. 2c**). Additionally, the majority of H3K27me3 peaks (76%) did not overlap with Kipf (**Extended Data Fig. 3c**). To determine whether one of the two marks, or both, could best predict piRNA clusters, we divided the genome into 1 kb bins and ranked these bins based on their level of histone mark enrichment. To enable comparisons between different marks, we calculated their classification performance as area under the ROC curve (AUC) (**Extended Data Fig. 3d**). We used published HP1a ChIP-seq data (Zenk et al. 2021) as a proxy for H3K9me3, as we noticed that the H3K9me3 antibody we used appeared to capture some H3K27me3 signal (see **Supplementary Note 2**). Based on analysis of the AUC, we found that H3K27me3 is an accurate predictor of dual-strand piRNA cluster location (AUC 0.867), similar to the predictive power of HP1a binding (AUC 0.842) but combining both marks further improved classification performance as a predictor of dual-strand piRNA clusters (AUC 0.927) (**Extended Data Fig. 3e**). Of note, we observed that H3K27me3 enrichment varied between piRNA source loci in agreement with E(z) dependence in piRNA production: clusters *42AB* and *38C* showing strong enrichment in H3K27me3 (**Fig. 2d,e**), whereas other clusters such as *20A* and *80F* whose piRNA production was not disrupted by E(z) depletion displayed lower levels of H3K27me3 (**Fig. 2d,f**). These observed correlations between H3K27me3/H3K9me3 co-occupancy and the production of piRNAs points towards a role for both H3K27me3 and H3K9me3 in the specification of piRNA cluster identity.

**Figure 3:**
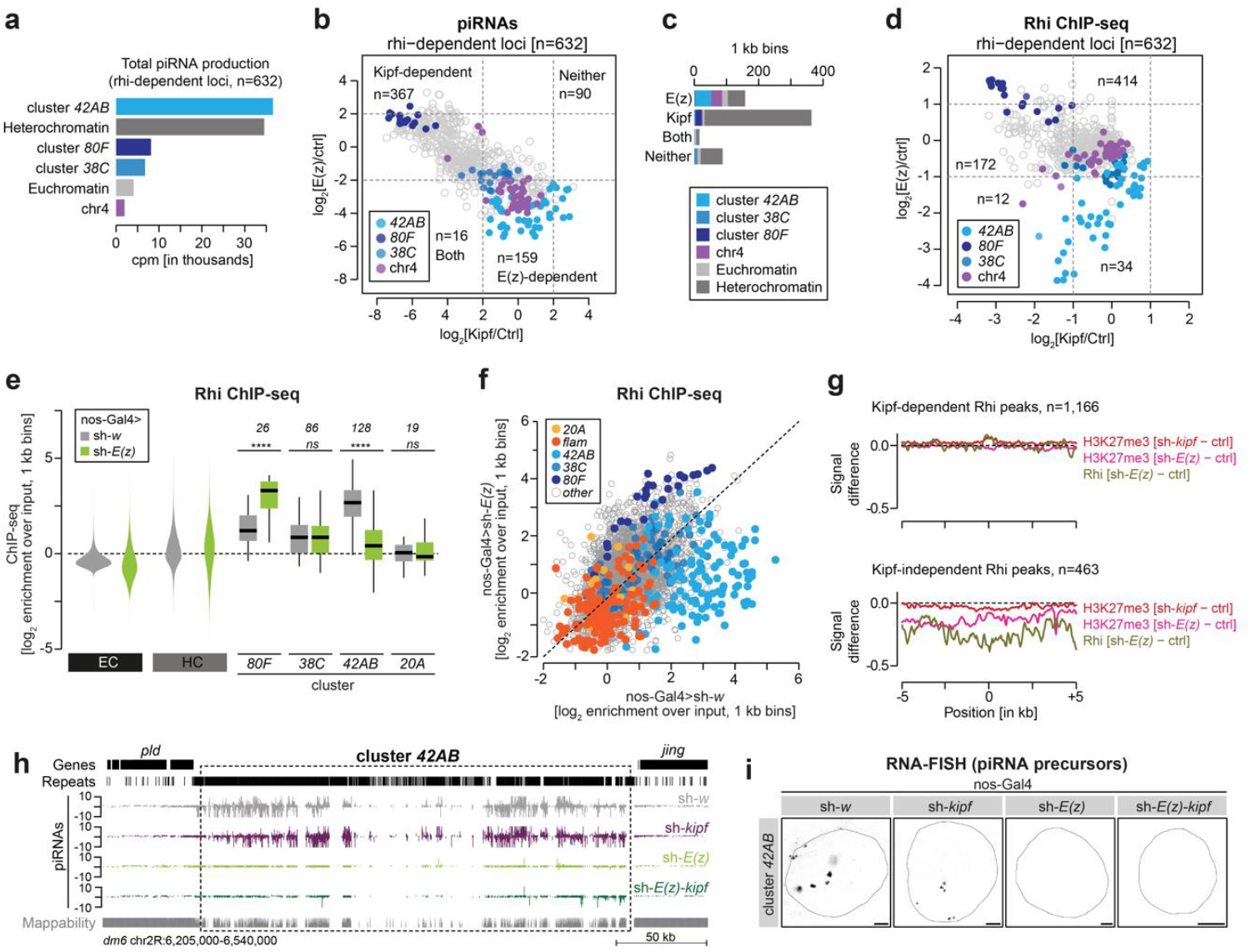
Rhi binding at Kipf-independent loci requires E(z) **a**, Bar graph showing genomic origin of Rhi-dependent piRNAs. The analysis was done across 1 kb bins (n=632) that displayed Rhi-dependency (>2-fold mean piRNA loss upon Rhi depletion) across several depletion strategies (pTOsk-Gal4 or MTD-Gal4 driven knockdown, and *rhi* knockout; 1-2 replicates each). **b**, Scatter plots showing log_2_-fold mean changes in levels of uniquely mapping piRNAs in 1 kb bins relative to control across several depletion strategies (nos-Gal4 or pTOsk-Gal4 driven knockdown of *E(z)*, 2 replicates each; MTD-Gal4 driven knockdown of *kipf* or *kipf* knockout, 1 replicate each). Only Rhi-dependent regions are shown as defined in a. Bins were categorized into E(z)*-*dependent and/or Kipf-dependent if displaying at least a 4-fold mean loss of piRNAs. **c**, Bar graph showing genomic origin for 1 kb bins per E(z)/Kipf-dependency category as defined in b. **d**, Scatter plot showing log_2_ fold change in Rhi ChIP-seq enrichment following depletion of E(z) using nos-Gal4 or Kipf using MTD-Gal4 across 632 Rhi-dependent piRNA loci as defined in a. Dashed lines indicated a 2-fold mean loss of Rhi occupancy. **e**, Violin plots (left) and box plots (right) showing average log_2_-fold Rhi enrichment by ChIP-seq from ovaries with nos-Gal4 driven *E(z)* knockdown versus control (average of 2 replicates each) in heterochromatin (HC) and euchromatic chromosome arms (EC), quantified across 1 kb bins (excluding piRNA clusters). Rhi occupancy at indicated piRNA clusters is shown as box plot quantification (n depicts 1 kb bins analysed for each piRNA cluster). **** corresponds to P<0.0001 based on Wilcoxon signed-rank test. Box plots show median (centre line), with interquartile range (box) and whiskers indicating at most 1.5x interquartile range). **f**, Scatter plot of genomic 1 kb bins contrasting average log_2_-fold ChIP-seq enrichment of Rhi in ovaries with nos-Gal4 driven *E(z)* knockdown versus control (average of 2 replicates each). **g**, Metaplot showing mean difference in H3K27me3 (pink) and Rhi (beige) ChIP-seq signal upon nos-Gal4-driven *E(z)* knockdown and H3K27me3 (red) ChIP-seq signal upon nos-Gal4-driven *kipf* knockdown across Rhi peaks categorised as either Kipf-dependent or not (see Methods). **h**, UCSC genome browser tracks of the region comprising dual-strand piRNA cluster *42AB* displaying uniquely mapping piRNAs (coverage normalized to miRNA reads) upon the indicated (double-)knockdowns as well as tracks indicating genes and repeats. **i**, Confocal images showing RNA-FISH signal for transcripts derived from dual-strand piRNA cluster *42AB* in control, *E(z*), and *kipf* knockdown as well as in *E(z)*-*kipf* double-knockdown, using the nos-Gal4 driver (scale bar: 5 μm).

### Rhi binding to a subset of piRNA source loci depends on the histone methyltransferase E(**z**)

The dual-strand cluster *42AB* is the major source of Rhi-dependent piRNAs in germ cells (**Fig. 3a**), and our data thus far support that its piRNA production is E(z)-dependent. Prior work implicates Kipf in guiding Rhi to a subset of piRNA source loci that includes piRNA cluster *80F*, which we find is E(z)-independent (**Fig. 1d-f, h, Extended Data Fig. 2d-f, h**), but not *42AB* (Baumgartner et al. 2022). We therefore hypothesized that Rhi binding to piRNA source loci that are independent of Kipf may depend on E(z). To test this hypothesis, we compared piRNA production from Rhi-dependent loci (n=632, ≥2-fold piRNA reduction after Rhi loss) in Kipf or E(z)-depleted ovaries. We observed that E(z), but not Kipf, is required for efficient piRNA production from cluster *42AB* and piRNA-producing loci on chr4, whereas cluster *80F* and most heterochromatic regions are E(z)-independent but require Kipf (**Fig. 3b,c**). Other piRNA clusters, such as *38C*, depend weakly on both Kipf and E(z) (**Fig. 3b**). Together, this suggests that E(z) and Kipf may provide two distinct modes of Rhi recruitment to separate subsets of piRNA source loci.

Production of piRNAs from dual-strand clusters in germ cells depends exclusively on Rhi recruitment and we therefore expect piRNA production patterns to closely reflect Rhi binding (Mohn et al. 2014). To determine whether E(z) directs Rhi binding to selected loci, we next performed Rhi ChIP-seq in E(z)-depleted ovaries and compared this to previously published Rhi ChIP-seq in Kipf-depleted ovaries (Baumgartner et al. 2022). Indeed, we observed that the Rhi binding pattern closely mirrored piRNA production (**Fig. 3d**). Consequently, we found that upon nos-Gal4-mediated E(z) depletion, Rhi enrichment was increased at Kipf-dependent piRNA source loci and reduced at those that depend on E(z) (**Extended Data Fig. 4a**). We next asked whether depletion of E(z) affects the global genomic distribution of Rhi. In agreement with the largely unchanged localization of Rhi by immunofluorescence (**Extended Data Fig. 2a**), we found that Rhi was not globally redistributed between euchromatin and heterochromatin upon knockdown of *E(z)* (**Fig. 3e**). Instead, we detected a strong reduction of Rhi binding at specific regions, including the *ey*/*Sox102F* region on chromosome 4 and the dual-strand piRNA cluster *42AB* (**Fig. 3e, Extended Data Fig. 4b**,**c**), whereas other loci, such as the *80F* piRNA cluster, accumulated more Rhi than in control ovaries (**Fig. 3e**). Those observations agreed with changes in H3K27me3 signal at these loci (**Extended Data Fig. 4d**). Notably, the observed changes in Rhi occupancy following E(z) depletion (**Fig. 3f**) followed the same pattern as piRNA precursor levels, which showed reduced levels for cluster *42AB* and an enrichment at cluster *80F* (**Fig. 1d-e**). Moreover, the observed differences in Rhi binding do not appear to be a consequence of changes in H3K9me3, as levels in this histone modification remain essentially unaltered across the genome (**Extended Data Fig. 4b**,**c**,**e**).

**Figure 4:**
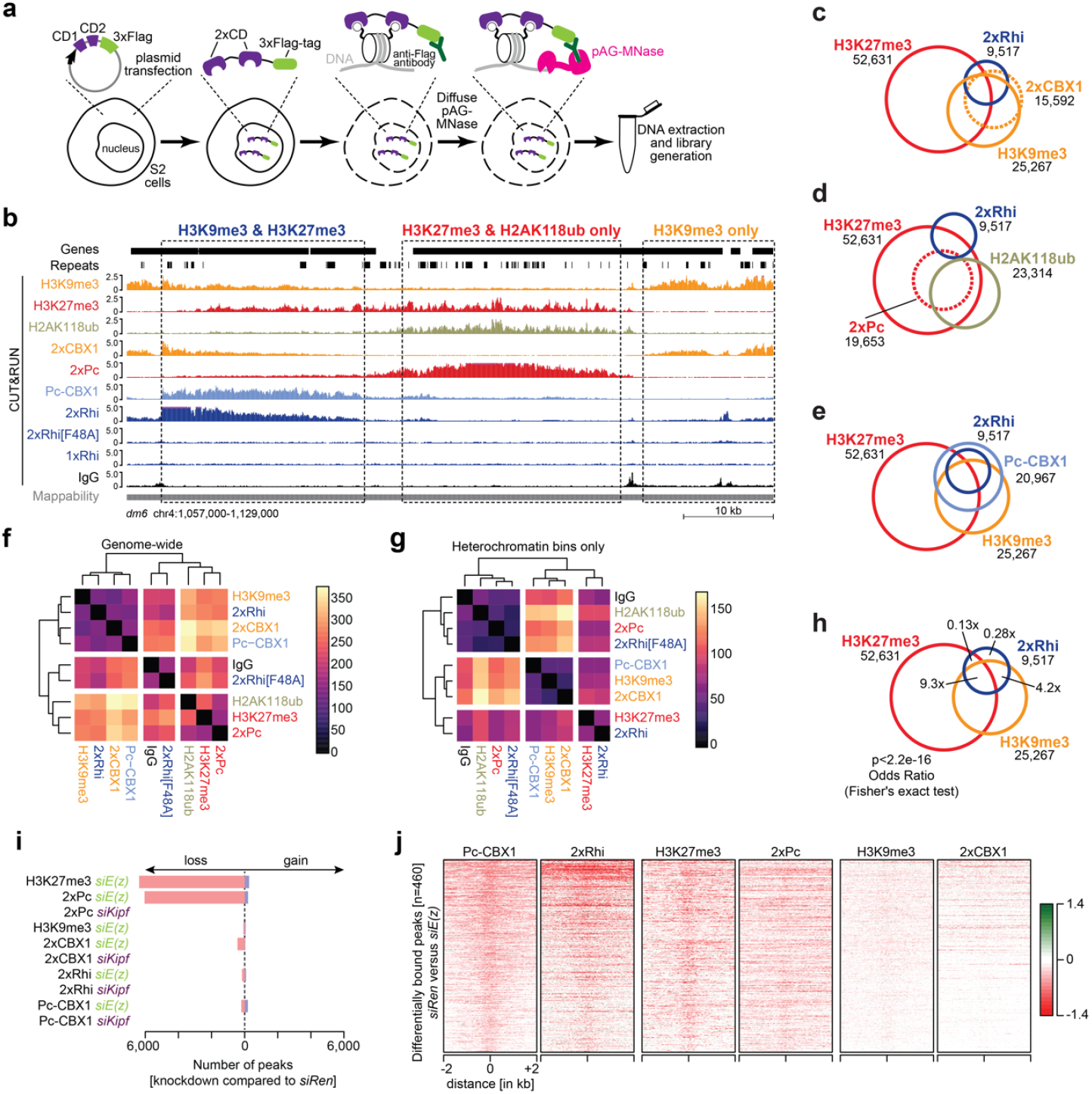
An *in vitro* system that recapitulates Kipf-independent Rhi binding. **a**, Schematic showing the experimental workflow for the chromodomain binding assay. **b**, UCSC genome browser tracks of a 72 kb region on chromosome 4 displaying the binding profiles of the indicated histone modifications and chromodomain constructs measured by CUT&RUN (counts per million [cpm] across pooled replicates). Dashed boxes indicate representative areas with either both H3K9me3 and H3K27me3, only H3K27me3, or only H3K9me3, respectively. The mappability of 100 nt reads is indicated below the tracks. **c**, Venn diagram showing overlap between CD^2xCBX1^ binding, H3K9me3, H3K27me3, and CD^2xRhi^ binding across 125,499 genomic 1 kb bins. Signal was considered to be present if the bin overlapped a MACS2 broad peak (q<0.05, broad-cutoff<0.1) present in at least two biological replicates. **d**, Same as c, but showing CD^2xRhi^ binding, H2AK118ub, H3K27me3, and CD^2xPc^ binding. **e**, Same as c, but showing CD^Pc-CBX1^ and CD^2xRhi^ binding, H3K27me3, and H3K9me3. **f**, Heatmap and hierarchical clustering (Euclidian distance) of CUT&RUN signal detected for the indicated chromodomain constructs and histone modifications [log_10_(fpkm)]. Signal shown as mean across biological replicates. Individual replicates shown in **Extended Data Fig. 6d. g**, Same as f, but across a subset of the genome classified as heterochromatin (10,180 out of 125,499 1 kb bins). Individual replicates shown in **Extended Data Fig. 6e. h**, Same as c, but showing H3K9me3, CD^2xRhi^ binding, and H3K27me3. Odds ratios for CD^2xRhi^ binding to H3K9me3, H3K27me3, both or neither were calculated using Fisher’s exact test. See **Extended Data Fig. 6f** for all major overlaps. **i**, Overview of differentially bound regions (called using DiffBind, padj<0.05) following *kipf* or *E(z)* knockdown. Consensus peaks (n=11,053) were called using DiffBind with MACS2 broad peaks from the indicated samples. **j**, Heatmaps showing the log_2_-fold change of *E(z)* knockdown compared to control knockdown for indicated constructs and histone modifications across Pc-CBX1 differentially bound regions.

We next analysed published ChIP-seq data (Baumgartner et al. 2022) for Rhi and Kipf occupancy across the genome. Based on the Rhi ChIP-seq, we identified 1,629 high-confidence Rhi binding sites using MACS2 (q<0.05, >4-fold enrichment). Out of these, 1,166 Rhi peaks were lost in *Kipf*-KD and were denoted as Kipf-dependent, while 463 peaks were retained and denoted as Kipf-independent (**Extended Data Fig. 4f**). In agreement with two separate modes of Rhi recruitment, we observed no reduction in H3K27me3 levels or Rhi occupancy at Kipf-dependent Rhi peaks following E(z) depletion, in line with these loci lacking H3K27me3 marks and suggesting that Rhi is recruited by Kipf in an H3K27me3-independent manner. In contrast, we observed a broad loss of both H3K27me3 and Rhi occupancy at Kipf-independent Rhi peaks upon E(z) depletion (**Fig. 3g**). In contrast, H3K27me3 was not affected by Kipf depletion (**Fig. 3g**). This suggests a strong correlation between the reduction in Rhi binding at Kipf-independent sites and the loss of H3K27me3 following *E(z)* knockdown.

Our polyA-selected RNA-seq data indicated an up to 6-fold reduction in mRNA levels of Moonshiner (Moon), a paralogue of TFIIA required for piRNA production from the majority of Rhi-dependent piRNA source loci (Andersen et al. 2017), in E(z) depleted ovaries (**Table S2**). We also observed reduced Rhi binding at Kipf-independent peaks in published ChIP-seq from *moon* KD (Andersen et al. 2017) (**Extended Data Fig. 4g**). It is thus a possibility that Rhi loss upon E(z) depletion may in part be driven by the reduced Moon levels. However, we consider secondary effects driven by Moon reduction unlikely since its loss was reported to lead to a complete loss of piRNAs from cluster *80F* and an increase in cluster *38C* derived piRNAs (Andersen et al. 2017), opposite to what is observed in our data. We therefore attribute the reduction in Rhi binding at Kipf-independent sites to loss of H3K27me3 upon *E(z)* knockdown.

Overall, our results provide evidence that H3K9me3 and H3K27me3 co-exist on a subset of piRNA source loci and appear to be a prerequisite for Rhi recruitment at these regions. Our results indicate that two non-redundant mechanisms influence Rhi binding to different piRNA source loci, namely E(z)-dependent and Kipf-dependent recruitment.

### Kipf and E(z) independently guide Rhi to its targets

To further investigate the idea of two non-redundant Rhi recruitment mechanisms, we next determined the effect of simultaneous knockdown of *E(z)* and *kipf* in *Drosophila* germ cells. We sequenced small RNAs from *E(z)*-*kipf* double-knockdown (dKD) ovaries and compared the piRNA levels to various control ovaries. First, when compared to knockdown of *w*, TE-derived sense and antisense piRNAs were reduced upon *E(z)* knockdown (**Extended Data Fig. 5a**). Notably, co-depletion of Kipf and E(z) resulted in more severe effects on piRNA levels (**Extended Data Fig. 5b**). However, in the absence of E(z), Kipf depletion has mild effects on piRNA populations (compare *E(z)-kipf* dKD with *E(z)-w* dKD; **Extended Data Fig. 5c**). In contrast, the combined depletion of E(z) and Kipf had pronounced effects on piRNA levels when compared to *kipf* knockdown alone (**Extended Data Fig. 5d**).

**Figure 5:**
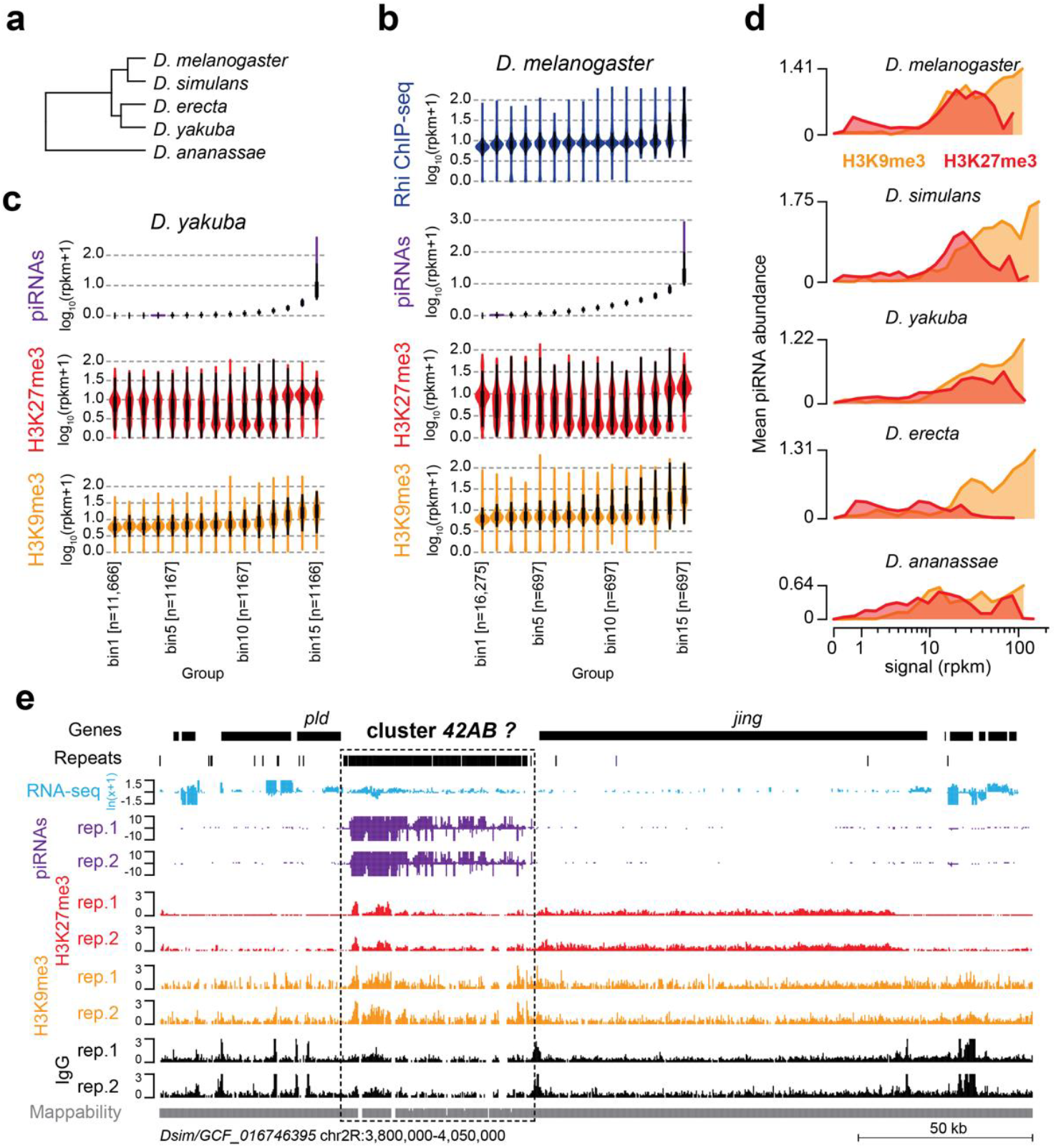
Conservation of Rhi association with H3K9me3 and H3K27me3 in Drosophilids. **a**, Phylogenetic tree of the five studied *Drosophila* species. **b**, Violin plots illustrating the distribution of piRNAs (purple), H3K9me3 (yellow) and H3K27me3 (red) CUT&RUN signal, and Rhi (blue) ChIP-seq levels [log_10_(rpkm+1)] in *D. melanogaster* ovaries. 10 kb bins across the genome were divided into 15 groups. Group 1 contains all 10 kb bins with no piRNA expression. The 14 remaining groups contain an equal number of 10 kb bins. Groups are ranked according to piRNA expression levels with the lowest level of the two strands shown to focus on dual-strand regions. **c**, Same as b, but showing piRNA, H3K9me3 and H3K27me3 levels [log_10_(rpkm+1)] for *Drosophila yakuba* ovaries. **d**, Line graphs showing mean piRNA abundance (n=3 replicates; signal shown as the lowest of the two strands, log_10_(rpkm+1)) plotted against H3K9me3 signal (orange; reads per kilo base per million mapped reads, rpkm) and H3K27me3 signal (red) for all 10 kb bins of the indicated *Drosophila* species (*D. melanogaster, D. simulans, D. yakuba, D. erecta*, and *D. ananassae)*. **e**, UCSC genome browser tracks from *D. simulans* displaying ribo-depleted RNA-seq (light blue), piRNAs (purple), and CUT&RUN for H3K27me3 (red) and H3K9me3 (yellow) as counts per million [cpm] at the *42AB*-syntenic region (dashed box). The mappability of 100 nt reads is indicated below the tracks.

Next, we checked piRNA levels at piRNA source loci. This revealed a strong reduction of piRNAs from the dual-strand cluster *42AB* in *E(z)-kipf* dKD and *E(z)* knockdown compared to *kipf* or control knockdowns (**Fig. 3h**). To quantify these observations, we performed a binning analysis and compared each (double)knockdown to respective controls (**Extended Data Fig. 5e**). As expected, piRNA production from the unistrand clusters *flam* and *20A* remained unchanged. Upon *E(z)-kipf* dKD we observed a 3.7-fold stronger reduction of piRNAs derived from cluster *42AB* compared to E(z) depletion alone (**Extended Data Fig. 5e-g**). As seen for TE-derived piRNAs, compared to E(z) depletion, the effect of *E(z)*-*kipf* dKD on cluster-mapping piRNAs was mild, while the opposite was true when compared to *kipf* knockdown alone (**Extended Data Fig. 5e**,**h**,**i**). Unexpectedly, while *kipf* knockdown alone resulted in reduced piRNA levels from cluster *80F*, we observed a recovery of piRNA production to near-normal levels in *E(z)-kipf* dKD ovaries (**Extended Data Fig. 5e**,**g**). These results suggest that in the absence of both E(z) and Kipf, Rhi nevertheless is able to bind to regions in this piRNA source locus.

Next, using RNA-FISH experiments, we found that, similar to *E(z)* knockdown, no signals for *42AB* were detected in the *E(z)*-*kipf* dKD ovaries (**Fig. 3i**). These results confirmed that transcription of *42AB* piRNA precursors relies on E(z) and is Kipf-independent. Notably, Rhi, which typically accumulates in large, continuous structures near the nuclear envelope at satellite regions in *kipf* knockdowns (Baumgartner et al. 2022), becomes redistributed throughout the nucleus in the *E(z)-kipf* dKD, resembling control and E(z) depleted ovaries (**Extended Data Fig. 5j**). Overall, these results highlight distinct, non-redundant mechanisms of Rhi recruitment at different piRNA clusters with some relying on *E(z)* and others on Kipf, but no evidence of interdependence between both recruitment mechanisms.

### Rhi binds a H3K9me3/H3K27me3 combinatorial code in a Kipf-independent context

To decisively investigate whether Rhi binding is driven by H3K9me3/H3K27me3 dual marked chromatin domains, we made use of S2 cells. To assess the role of the combinatorial histone marks in this system, which lacks an endogenous piRNA pathway (Eastwood et al. 2021), we took advantage of well-characterized chromatin reader domains with known binding specificities. Recent work showed that synthetic dual-chromatin reader domains for specific histone marks can be used to detect chromatin modifications, including combinatorial histone codes, through the use of different binders linked in tandem (Villasenor et al. 2020). Rhi, like other HP1 family proteins, contains a chromodomain (CD) that binds H3K9me3-modified histone tails at its N-terminus and a chromoshadow domain at the C-terminal region, connected together via the hinge region (Vermaak et al. 2005; Le Thomas et al. 2014; Mohn et al. 2014; Yu and Huang 2015). Using CUT&RUN in S2 cells, we characterised the binding patterns of dual-CD constructs similar in design to Villasenor and colleagues (see Methods for details). Specifically, we generated dual-CD constructs to recognize H3K9me3 (CD^2xCBX1^), H3K27me3 (CD^2xPc^), or dually H3K9me3/H3K27me3-decorated domains (a bivalent chromodomain fusion CD^Pc-CBX1^) (**Fig. 4a, Extended Data Fig. 6a**). We confirmed that CD^2xCBX1^ bound to regions decorated in H3K9me3 (**Fig. 4b,c**), the CD^2xPc^ associated with regions enriched in H3K27me3 (**Fig. 4b**,**d, Extended Data Fig. 6b**), and the bivalent CD^Pc-CBX1^ reader exhibited preferential enrichment at loci marked by both H3K9me3 and H3K27me3 modifications (**Fig. 4b,e**).

Next, we tested whether the association of tandem Rhi chromodomains (CD^2xRhi^) recapitulated the binding patterns observed for the bivalent H3K9me3/H3K27me3 reader CD^Pc-CBX1^ (**Fig. 4b,e**). Strikingly, while CD^2xRhi^ binding strongly resembled that of CD^2xCBX1^ and the bivalent CD^Pc-CBX1^ genome-wide (**Fig. 4f, Extended Data Fig. 6c**,**d**), within heterochromatic regions, CD^2xRhi^ binding instead strongly correlated with H3K27me3 levels (**Fig. 4g, Extended Data Fig. 6e**). In contrast, Rhi dual-chromodomains containing a F48A point mutation in the aromatic cage (CD^2xRhi[F48A]^) essentially recapitulated the IgG control (**Fig. 4b,f-g**). Although CD^2xRhi^ binding was strong already at loci with only H3K9me3 (Odds Ratio [OR] 4.2, Fisher’s exact test), the strongest CD^2xRhi^ binding was observed at loci marked with both H3K9me3 and H3K27me3 (OR 9.3), whereas binding was depleted at loci only marked with H3K27me3 (OR 0.13) (**Fig. 4h**). Of note, regions marked by both H3K9me3 and H3K27me3 showed no presence of the PRC1 deposited mark H2AK118ub generally associated with the presence of H3K27me3 (Margueron and Reinberg 2011) (**Fig. 4b**). These results indicate that Rhi recruitment depends on H3K27me3 but is independent of H2AK118ub (**Fig. 4b,d**). Indeed, a binning analysis revealed that CD^2xRhi^ binding in S2 cells clearly differs from that of CD^2xCBX1^ and CD^2xPc^ through its ability to bind both clusters *80F* and *42AB*, which are strong CD^2xCBX1^ targets, and the CD^2xPc^-targeted *38C* region (**Extended Data Fig. 6g**,**h**). Taken together, these results

show that, similar to CD^Pc-CBX1^, CD^2xRhi^ preferentially binds to regions decorated with both H3K9me3 and H3K27me3. To test whether CD^2xRhi^ binding in S2 cells was mediated by Kipf, we repeated the CD experiments upon depletion of Kipf. Despite robust knockdown of *kipf* (**Extended Data Fig. 6i**) and construct expression (**Extended Data Fig. 6j**), we did not observe differences in the binding patterns of any CD construct (**Fig. 4i, Extended Data Fig. 6k-n**). Importantly, upon *kipf* knockdown, CD^2xRhi^ binding was indistinguishable from the control knockdown (**Extended Data Fig. 6n**), hence suggesting that binding by the Rhi chromodomain in this assay is Kipf-independent.

### Kipf-independent Rhi binding depends on E(**z**)

To directly test the importance of H3K27me3 in specifying Rhi chromodomain binding, we performed CUT&RUN in S2 cells for CD^2xCBX1^, CD^2xPc^, CD^Pc-CBX1^, and CD^2xRhi^ upon control [*siRen*] or E(z) knockdown [*siE(z)*]. Principle component analysis (PCA) showed high levels of similarities between genome-wide CD^2xPc^ binding and H3K27me3 distribution and between genome-wide H3K9me3 distribution and CD^2xCBX1^ binding (**Fig. 4i, Extended Data Fig. 7a**). PCA also confirmed, as we observed in earlier experiments (**Fig. 4b, Extended Data Fig. 6c**), that the binding pattern of CD^2xRhi^ in control S2 cells more closely resembles that of the CD^Pc-CBX1^ binding pattern over that of CD^2xCBX1^ or CD^2xPc^ (**Extended Data Fig. 7a**). In the *E(z)* knockdown cells, binding of CD^2xPc^ was strongly affected while CD^2xCBX1^ and H3K9me3 distribution remained mostly unchanged (**Fig. 4i**). Notably, both CD^2xRhi^ and CD^Pc-CBX1^ distribution become more similar to that of CD^2xCBX1^ and H3K9me3 in the absence of *E(z)*, consistent with a reduced importance of H3K27me3 for their distribution (**Extended Data Fig. 7a**).

Next, to further determine the impact of H3K27me3 on Rhi binding, we divided the genome onto 1 kb bins. Bins were categorised into six different groups based on the degree of H3K27me3 loss upon depletion of E(z). We then plotted for each group the fold-change in binding levels of the different chromodomains as assessed by CUT&RUN in *siE(z)* compared to *siRen*. While loss of H3K27me3 strongly reduced CD^2xPc^ binding, no major genome-wide effects were seen for CD^2xCBX1^, CD^Pc-CBX1^ or CD^2xRhi^ (**Extended Data Fig. 7b**, upper panels). This reflects the ability of CD^2xPc^ to bind all H3K27me3-decorated regions genome-wide, whereas only a subset of H3K27me3-decorated regions, those also marked with H3K9me3 can be bound by CD^Pc-CBX1^ and CD^2xRhi^. To identify H3K9me3 and H3K27me3 dual marked regions affected by the loss of E(z), we focused on CD^2xCBX1^-bound regions (as a proxy for H3K9me3) and observed that depletion of E(z) resulted in reduced binding of both CD^2xRhi^ and CD^Pc-CBX1^ in these H3K9me3-enriched bins (**Extended Data Fig. 7b**, lower panels). As expected, we observed no differences when performing the same analysis for *siKipf* compared to control knockdown (**Extended Data Fig. 7c**).

We next identified 460 CD^Pc-CBX1^ binding sites where its binding was significantly reduced (DiffBind, padj<0.05) after H3K27me3 loss upon E(z) depletion. Notably, these regions displayed a similar loss of CD^2xRhi^ binding following H3K27me3 depletion, despite retaining similar H3K9me3 levels (**Fig. 4j, Extended Data Fig. 7d**). These results indicate that binding of CD^Pc-CBX1^ and CD^2xRhi^ at these H3K9me3/H3K27me3-decorated loci require the presence of both H3K27me3 and H3K9me3 marks, and that H3K9me3 alone is not sufficient for binding. Taken together, these findings indicate that a combinatorial H3K9me3/H3K27me3 histone code, that is independent of Kipf but recognized by the Rhi chromodomain, underlies Rhi binding specificity at a subset of piRNA source loci.

### The combinatorial histone code H3K9me3/H3K27me3 at major piRNA source loci is conserved in Drosophilids

It has previously been shown by Rhi ChIP-seq that the *Drosophila simulans* Rhi CD has similar binding specificity as those of *Drosophila melanogaster* when expressed in *D. melanogaster* ovaries (Parhad et al. 2017). Based on this, we hypothesized that Rhi association with H3K9me3/H3K27me3 dual domains could be conserved across species. This is further supported by the high sequence conservation of the Rhi CD across *Drosophila* species (Vermaak et al. 2005). In particular, aromatic residues in the CD, important for binding to methylated lysine residues, show high conservation (**Extended Data Fig. 8a**), suggesting that Rhi chromodomains in Drosophilids all have the potential to bind H3K9 and H3K27 tri-methylated histone tails.

To test this, we assessed by CUT&RUN the presence of H3K9me3 and H3K27me3 at piRNA producing loci in ovaries from *Drosophila simulans* (MRCA 5.6 MYA), *Drosophila erecta* (MRCA 8.2 MYA), *Drosophila yakuba* (MRCA 11.4 MYA), and *Drosophila ananassae* (MRCA 33.9 MYA) (**Fig. 5a**). We compared our CUT&RUN data for H3K9me3 and H3K27me3 marks with small RNA-seq of ovaries from these species (van Lopik et al. 2023). We divided the genomes into 10 kb bins that were then classified onto 15 groups according to piRNA levels present in each species. We observed that at least for *Drosophila melanogaster* and *Drosophila yakuba*, bins with high levels of piRNAs also showed strong signals of H3K9me3 and H3K27me3 marks, whereas the absence or low levels of piRNAs correlated mostly with the presence of H3K27me3 and absence of H3K9me3 (**Fig. 5b**,**c, Extended Data Fig. 8b**). Genome-wide analyses showed that piRNA producing loci are preferentially decorated with both H3K9me3 and H3K27me3 in *Drosophila simulans, Drosophila yakuba, Drosophila erecta* and *Drosophila ananassae* (**Fig. 5d**). We also detected both histone marks at the region of the *Drosophila simulans* genome that is syntenic to the *Drosophila melanogaster 42AB* locus (**Fig. 5e**). These results suggest that the combination of H3K9me3 and H3K27me3 in regions where Rhi binds is a conserved feature in Drosophilids.

In conclusion, our study identifies the H3K27 methyltransferase E(z) as an important regulator of TE expression in *Drosophila* germ cells. We show that, in addition to the well-described H3K9me3 mark, H3K27me3 is important for guiding Rhi binding in a Kipf-independent manner, thus contributing to the definition of a dual-strand piRNA source loci.

## DISCUSSION

Chromatin structure and gene expression are strongly influenced by complex patterns of histone modifications, such as methylation, acetylation, phosphorylation, and ubiquitination. It is becoming increasingly clear that these modifications occur simultaneously, creating a diverse range of potential combinations, also referred to as the combinatorial histone code, and these patterns can be interpreted by various reader/effector proteins.

Here we provide evidence that the co-occurrence of the histone modifications H3K9me3 and H3K27me3, usually thought to occupy distinct domains, plays an important role in marking genomic loci for specific binding of the heterochromatin protein Rhi, a key step to promote piRNA source loci expression. By performing a screen for novel factors important for TE control, we identified the H3K27me3 methyltransferase E(z) and demonstrated that several Rhi-bound loci that produce vast amounts of piRNAs in germ cells are decorated with both H3K9me3 and H3K27me3 marks. We demonstrate that at these loci, Rhi recruitment is independent of the zinc finger protein Kipf and instead requires the H3K27me3 methyltransferase E(z), and that Rhi likely binds H3K27me3 and H3K9me3 through its chromodomain. These data demonstrate the existence of two independent modes of Rhi recruitment at distinct subsets of piRNA source loci: that reliant on Kipf-recognized DNA sequence and that dependent on chromatin context as established by E(z). Our results also point to the involvement of additional complexes in TE regulation, such as the COMPASS and NSL complex. Of note, a recent study has shown that the depletion of germline-specific subunits of the NSL complex (NSL1, NSL2, and NSL3) resulted in reduced piRNA production from telomeric piRNA source loci, thus emphasizing the role of the NSL complex in piRNA precursor transcription within these loci (Iyer et al. 2023).

This work is the first report and functional exploration of the co-occupancy of H3K9me3 and H3K27me3 modifications at piRNA source loci in *Drosophila*. While H3K9me3 and H3K27me3 histone marks have previously been suggested to be mutually exclusive, our work and that of others demonstrate that they co-exist in a number of contexts, although until now without a clear function (Liu et al. 2011; Ho et al. 2014; Mozzetta et al. 2014; Evans et al. 2016; Walter et al. 2016; Ichihara et al. 2022). Indeed, the co-occurrence of H3K9me3 and H3K27me3 at TEs has been reported in other species. In the ciliate *Paramecium tetraurelia* and in mice the deposition of both marks play a crucial role in maintaining control over a wide range of TEs, thereby ensuring the long-term stability of the genome (Walter et al. 2016; Frapporti et al. 2019). The H3K27me3 mark was proposed to act as a backup system for TE silencing in low DNA methylation contexts in flowering plants and in mammals (Mathieu et al. 2005; Weinhofer et al. 2010; Deleris et al. 2012; Saksouk et al. 2014; Walter et al. 2016; Rougee et al. 2021). In the ancestral eukaryotic lineage, such as the marine diatom *Phaeodactylum*, H3K27me3 is also enriched at TE sequences with or without co-occurring H3K9me3 (Veluchamy et al. 2015; Deleris et al. 2021). Interestingly, *C. elegans* piRNA source loci that give rise to 21U-RNAs are found within H3K27me3-rich genomic regions (Beltran et al. 2019). In addition, recruitment of the *C. elegans* Upstream Sequence Transcription Complex, required for piRNA precursor transcription, is guided by the chromodomain-containing protein UAD-2 which can bind H3K27me3 (Huang et al. 2021). In the absence of H3K27me3, UAD-2 fails to be recruited to piRNA source loci, resembling the effect observed for Rhi upon germline depletion of E(z).

Although E(z)-catalysed H3K27me3 appears to be required for Rhi recruitment to a subset of piRNA source loci, the mechanisms by which E(z) is recruited and when H3K27me3 is deposited at specific *Drosophila* dual-strand piRNA source loci during development remains to be determined. Intriguingly, in *Paramecium*, the Polycomb protein Ezl1 plays an effector function and is targeted directly to TE insertions via interaction with a PIWI protein and subsequently mediates H3K9me3 and H3K27me3 deposition at these loci (Miro-Pina et al. 2022). *Drosophila* E(z) has been shown to co-immunoprecipitate with Piwi in ovaries (Peng et al. 2016), which hints at a possible role of PIWI proteins in guiding E(z) to deposit H3K27me3 at piRNA source loci. Moreover, the presence of H3K27me3 at these loci in multiple cell types, including germ cells and cells derived from the somatic lineage (e.g., S2 cells that are derived from *Drosophila* embryos), suggests that H3K27me3 is established early during development or possibly deposited maternally (Zenk et al. 2017), and retained somatically as development progresses. A previous study demonstrated divergence in the spatio-temporal expression patterns of Kipf and Rhi, with Kipf exhibiting very low levels in ovarian germline stem cells and cystoblasts (Baumgartner et al. 2022). In light of this observation, we suggest that H3K27me3 may collaborate with H3K9me3 during the early stages of oogenesis, where Rhi functions independently of Kipf. Understanding how and when histone methyltransferases are guided to piRNA source loci and their other targets at which they deposit histone marks is a critical further question to better understand the piRNA system and its interplay with chromatin biology.

While bivalent chromatin, characterised by the presence of both H3K4me3 and H3K27me3, or dual domains have been previously suggested to play important roles in regulating gene expression, the diversity of chromatin reader domains able to interpret either each histone post-translational modification or exhibiting dual recognition of both modifications *in vitro* brings additional complexity to the understanding of chromatin biology (Azuara et al. 2006; Mikkelsen et al. 2007; Du et al. 2012; Gaertner et al. 2012; Musselman et al. 2012; Kumar et al. 2021; Mashtalir et al. 2021; Barral et al. 2022). Proteins with a paired domain, such as Trim24, which contains a PHD and a Bromo domain targeting unmodified H3K4 and acetylated H3K23, respectively, has been shown to bind a single histone tail bearing these two marks *in vitro* (Tsai et al. 2010). However recent work indicates that H3K23ac is absent from Trim24-binding sites in mouse embryonic stem cells and does not appear to play a role in Trim24 localization *in vivo* (Isbel et al. 2023).

Here, we showed that tandem Rhi chromodomains have a similar binding pattern as bivalent chromodomain fusion proteins expected to bind H3K9me3/H3K27me3 decorated domains, strongly suggesting that the Rhi chromodomain binds simultaneously to both H3K9me3/H3K27me3. Unlike the chromodomain of HP1a, the Rhi CD has been reported to form a homodimer in two crystallographic studies (Le Thomas et al. 2014; Yu et al. 2015). To investigate whether Rhi chromodomain dimers have the capacity to bind a histone H3 tail carrying dual-modified K9me3/K27me3, we successfully modelled the binding of the Rhi CD dimer to a single 40 amino acid histone H3 tail peptide carrying simultaneous tri-methylation of lysine 9 and lysine 27 using molecular dynamics (MD) simulations (**Extended Data Fig. 8c**). However, more recent work using an orthogonal approach (size-exclusion chromatography with inline multi-angle light scattering) failed to detect Rhi dimerization in solution (Baumgartner et al. 2024). Moreover, despite detecting robust expression (**Extended Data Fig. 6j**), we were unable to obtain specific binding patterns when performing profiling experiments in S2 cells using a construct carrying a single Rhi CD (CD^1xRhi^) (**Extended Data Fig. 8d**), possibly arguing against the ability of this construct to dimerise *in vivo*. However, other reports suggested that the binding affinity of individual CD constructs are insufficient for efficient chromatin profiling (Villasenor et al. 2020), hence complicating the interpretation of these results. While our work suggests a role for a dual histone code in determining Rhi binding, likely in a CD-dependent manner, we cannot exclude alternative mechanisms involving an additional co-factor that associates with H3K27me3 or E(z). Considering that previous work found that the Rhi CD does not bind H3K27me3 alone *in vitro* (Mohn et al. 2014; Yu et al. 2015), one interesting hypothesis is the presence of a yet to be discovered co-factor that interacts with the Rhi chromodomain and H3K27me3. Considering that E(z) was recently identified as potential interactor of Del in IP-MS data from *D. simulans, D. erec*ta and *D. virilis* (Riedelbauch et al. 2024), another hypothesis could be that in *D. melanogaster* this interaction is mediated by the Rhi chromodomain directly. Alternatively, the presence of both H3K9me3 and H3K27me3 could provide a chromatin context (e.g., chromatin compaction and/or repression of canonical transcription) more favourable to Rhi binding compared to the presence of H3K9me3 alone.

In summary, our study provides an unexpected example highlighting *in vivo* the importance of a combinatorial histone code in the binding specificity of a chromatin binding protein.

## MATERIALS AND METHODS

### Data generation

#### Fly husbandry and stocks

All flies were kept at 18 °C or 25 °C on standard cornmeal or propionic food. Flies were obtained from the Vienna *Drosophila* Resource Center (VDRC), or from the Bloomington *Drosophila* Stock Center (BDSC). All used fly stocks are listed in **Table S3**. For germline-specific knockdowns, we used a fly line containing a UAS-Dcr2 transgene and a nos-GAL4 driver (Czech et al. 2013), a pTOsk-GAL4 driver (ElMaghraby et al. 2022), or a nos-GAL4 driver (BDSC:4937), each crossed to stocks expressing shRNAs (Ni et al. 2011) or dsRNAs (Perkins et al. 2015) under the UAS promoter. After mating at 27 °C for five days, parental flies were removed from the vials. Hatching F1 offspring were collected and aged with yeast for 2-3 days prior to use for follow-up experiments.

#### RNA isolation and qRT-PCR

Samples were lysed in 1 ml TRIzol reagent (Thermo Fisher Scientific) and RNA was extracted according to manufacturer’s instructions. One microgram of total RNA was treated with DNaseI (Thermo Fisher Scientific), and reverse transcribed with the Superscript III First Strand Synthesis Kit (Thermo Fisher Scientific), using oligo(dT)20 primers. Primer sequences are listed in **Table S4**.

#### TE screen

One to two different fly lines per gene (1 replicate per line) expressing shRNAs or dsRNAs under the UAS promoter (Ni et al. 2011; Perkins et al. 2015) were crossed with a fly line containing a UAS-Dcr2 transgene and a nos-GAL4 driver as previously described (Czech et al. 2013). RNA was isolated using 5-10 ovaries per cross and reverse transcribed as described above. Multiplexed qPCRs were carried out using TaqMan Universal Master Mix II, no UNG (Applied Biosystems) as previously described (Czech et al. 2013). Experiments were performed on a CFX96 Real-Time System C1000 Touch Thermal Cycler (BioRad). Z-scores for TE expression were calculated on ΔCT values [CT(TE) – CT(*rp49* control)] (Livak and Schmittgen 2001). The primers and probes used are listed in **Table S4**. Positive hits were confirmed by qPCRs using SYBR green Master mix (Thermo Fisher Scientific). RT-qPCRs were performed on a QuantStudio Real-Time PCR Light Cycler (Thermo Fisher Scientific).

#### RNA-FISH

Single-molecule RNA fluorescence in situ hybridization (RNA-FISH) for transcripts derived from piRNA clusters *42AB, 80F* and *20A*, as well as the TEs *Burdock, Gypsy12*, and *copia* was performed using Stellaris probes (Biosearch Technologies). Probe sequences are listed in **Table S4**. Ovaries from 3 to 6-day-old flies were dissected in Schneider’s *Drosophila* Medium and fixed in Fixing Buffer (4% formaldehyde, 0.3% Triton X-100, 1x PBS) for 20 min at room temperature, rinsed three times in 0.3% Triton X-100, once in PBS, and permeabilized in 70% ethanol at 4°C overnight. Permeabilized ovaries were rehydrated in RNA-FISH wash buffer (10% formamide in 2x SSC) for 10 min. Ovaries were resuspended in 50 μL hybridization buffer (10% dextran sulphate, 10% formamide in 2xSSC) supplemented with 1.5 μL of RNA-FISH probes. Hybridization was performed with rotation at 37°C overnight. Ovaries were then washed twice with RNA-FISH wash buffer at 37°C for 30 min and twice with 2xSSC solution. Then, DNA was stained with DAPI (1/500 dilution in 2x SSC) at room temperature for 20 min. Ovaries were mounted in 30 μL Vectashield mounting medium and imaged on a Zeiss Zeiss LSM-800 confocal microscope. The resulting images were processed using FIJI/ImageJ.

#### Immunofluorescence

Fly ovaries were dissected in ice-cold PBS, fixed for 14 min in 4% PFA (Alfa Aesar) at RT and permeabilized with 3×10 min washes in PBS with 0.3% Triton (PBS-Tr). Samples were blocked in PBS-Tr with 1% BSA for 2 hrs at RT and incubated overnight at 4 °C with primary antibodies in PBS-Tr and 1% BSA. After 3×10 min washes at RT in PBS-Tr, secondary antibodies were incubated overnight at 4 °C in PBS-Tr and 1% BSA. After 4x10min washes in PBS-Tr at RT with 4′,6-diamidino-2-phenylindole (DAPI; Invitrogen) added during the third wash, and 2x5 min washes in PBS, samples were mounted with ProLong Diamond Antifade Mountant (Thermo Fisher Scientific) and imaged on a Leica SP8 or Zeiss LSM-800 confocal microscope. Images were deconvoluted using Huygens Professional or using FIJI/ImageJ. All used antibodies are listed in **Table S5**.

#### Molecular cloning and constructs

All constructs were cloned using the NEBuilder HiFi DNA Assembly kit (New England Biolabs E2621) according to manufacturer’s instructions. Chromodomain (CD) sequences were amplified from cDNA prepared from ovaries or ordered as gBlock DNA fragments from Integrated DNA Technologies (IDT). The final constructs expressed the CDs of interest tagged amino-terminally with an NLS-3xFlag-EGFP cassette under the control of the *D. simulans ubiquitin* promoter. A list and link to the sequence of all constructs used in this study are provided in **Table S6**.

#### mRNA-seq (**polyA selected**)

Total RNA was extracted from 30 ovaries from 3-6 day-old flies using TRIzol (Thermo Fisher Scientific) in three replicates. One microgram of total RNA was subjected to polyA selection and subsequent fragmentation, reverse transcription, and library preparation according to the manufacturer’s instructions using the Illumina stranded mRNA Prep for sequencing. Sequencing was performed by Novogene on an Illumina Novaseq 6000 instrument.

#### ribo-depleted RNA-seq

Total RNA was extracted from 10 to 20 ovaries from 3-6 day-old flies using TRizol (Thermo Fisher Scientific) following the manufacturer’s instructions. For all the experiments conducted with the pTOsk-Gal4 driver, ribosomal RNA was depleted using RiboPOOL (siTOOLs, Biotech) as described (Munafo et al. 2021). RNA-seq libraries were produced using NEBNext Ultra Directional Library Prep Kit for Illumina, following the manufacturer’s instructions for rRNA depleted RNA. Library size distribution was analysed on a TapeStation instrument (Agilent Technologies) using a High Sensitivity D1000 ScreenTape. Libraries were pooled in equal molar ratio and quantified with KAPA Library Quantification Kit for Illumina (Kapa Biosystems) and sequenced paired-end 50 on an Illumina NovaSeq 6000 instrument. For the experiments conducted using the nos-Gal4d driver, rRNA depletion was performed from 1 µg total RNA using the RNA Depletion stranded Library Prep kit (BGI). The samples were then sequenced as PE100 reads on the DNBSEQ G400 sequencer, and adapter-clipped reads were provided by BGI.

#### Small RNA-seq

Small RNA extraction was performed as described (Grentzinger et al. 2020; van Lopik et al. 2023). Argonaute-bound small RNAs were isolated from 10 to 30 pairs of ovaries from 3-5 day-old flies using TraPR ion exchange spin columns (Lexogen, catalogue nr. 128.08). Small RNA libraries were generated using the Small RNA-seq Library Prep Kit (Lexogen; catalogue nr. 052.96) according to the manufacturer’s instructions. For all experiments conducted with the pTOsk-Gal4 driver, ovaries of the five *Drosophila* species, and dKD or *kipf* knockdown experiments using the nos-Gal4 driver, sequencing was performed at the CRUK CI Genomics core on an Illumina NovaSeq 6000 instrument. Sequencing for the remaining experiments conducted using the nos-Gal4 driver was performed by Fasteris SA (Geneva, Switzerland) on an Illumina NextSeq550 instrument.

#### ChIP-seq

Chromatin immunoprecipitation (ChIP) was performed as previously described in (Lee et al. 2006) with minor modifications. Briefly, 100 ovary pairs were manually dissected into Schneider media and cross-linked in 1% formaldehyde/PBS for 10 min at room temperature with agitation. The cross-linking reaction was quenched by STOP buffer (PBS 1X, Triton 0.1%, Glycine 1 M) and ovaries were washed in PBS and homogenized in a glass douncer: first slightly dounced in PBST 0.1% and centrifugated 1 min 400 g, followed by strong douncing in cell lysis buffer (KCL 85 mM, HEPES 5 mM, NP-40 0.5%, Sodium butyrate 10 mM, EDTA free protease inhibitor cocktail Sigma) following by 5 min centrifugation at 2000 g. We performed 2 washes with cell lysis buffer. The homogenates were then lysed on ice for 30 min in nucleus lysis buffer (HEPES 50 mM, EDTA 10 mM, N lauryl sarkosyl 0.5%, sodium butyrate 10 mM, EDTA free protease inhibitor cocktail Sigma). DNA was sheared using a Bioruptor pico from Diagenode for 10 cycles (30 sec on, 30 sec off). The sonicated lysates were cleared by centrifugation and then incubated overnight at 4°C with 5 µl of specific antibodies (**Table S5**). Then 40 µL of Protein A Dynabeads was then added and allowed to bind antibody complexes by incubation for 1 hr at 4°C. Following four washing steps with high salt buffer (Tris pH 7.5 50 mM, NaCl 500 mM, Triton 0.25%, NP-40 0.5%, BSA 0.5%, EDTA pH 7.5 5 mM), DNA-protein complexes were eluted and de-cross-linked 10 h at 65°C. RNA and protein was digested by RNase A and Proteinase K treatments, respectively, before purification using Phenol:Chloroform:Isoamyl Alcohol 25:24:1 (Sigma) according to the manufacturer’s instructions. Barcoded libraries were prepared using Illumina technology, and subsequently sequenced on a NextSeq High (Illumina) by Novogene (Rhi and H3K9me3 ChIP-seq), or by the Jean Perrin facility (H3K27me3 ChIP-seq).

#### CUT&RUN

1 million cells per sample were harvested and washed three times with Wash buffer (20 mM HEPES, pH7.5, 150 mM NaCl, 0.5 mM spermidine supplemented with protease inhibitors) and resuspended in 1 ml of Wash buffer. 10-20 fly ovaries were dissected in ice cold PBS per sample. Ovaries were digested in 250 µl of dissociation buffer (0.5% Trypsin and 2.5 mg/ml Collagenase A in PBS) for an hour with shaking at 800 rpm at 30 °C. The digestion was stopped with 250 µl of Schneider medium containing 10% FBS. Cell suspensions were filtered through 40 µm strainers, pelleted and washed three times with Wash buffer and resuspended in 1 ml of Wash buffer. Following sample preparation, CUT&RUN was performed according to instructions in CUT&RUN protocol V.3 with some modifications (Meers et al. 2019). 10 µl of activated Concanavalin A-coated magnetic beads (Bangs Laboratories) were added to each sample and rotated for 10 min at RT. Bead-bound cells were incubated with 5 µl of antibody (**Table S5**) and 95 µl of antibody buffer (20 mM HEPES pH 7.5, 150 mM NaCl, 0.5 mM spermidine, 0.05% digitonin) overnight at 4 °C. Bead-bound cells were washed twice with Dig-wash buffer (0.05% digitonin in Wash buffer). Bead-bound cells were resuspended in Dig-wash buffer containing 1x CUTANA pAG-MNase (Epicypher) and rotated for 1 hr at RT. Following pAG-MNase binding, bead-bound cells were washed twice with Dig-wash buffer and resuspended in 100 µl Dig-wash buffer. Chromatin digestion was performed on ice for 30 min by adding 2 μL of 100 mM CaCl_2_. Digestion was stopped by the addition of 2x STOP Buffer (340 mM NaCl, 20 mM EDTA, 4 mM EGTA, 0.05% digitonin, 100 µg/ml RNase A (Thermo Fisher Scientific), 50 µg/ml glycogen) and samples were incubated at 37 °C for 30 min to release DNA fragments into the solution. After centrifugation, 0.1% SDS and 0.2 μg/μl Proteinase K were added to the supernatant and samples were incubated for 1 hr at 50 °C. DNA was extracted using Phenol:Chloroform:Isoamyl Alcohol 25:24:1 (Sigma) according to the manufacturer’s instructions. Libraries were prepared following the manufacturer’s instructions with NEBNext Ultra II DNA Library Prep Kit for Illumina. Sequencing was performed on a NovaSeq 6000 instrument (Illumina).

#### Tissue culture, transfection and knockdowns

*Drosophila* Schneider 2 (S2) cells (Thermo Fisher Scientific; R69007) were cultured at 26 °C in Schneider’s *Drosophila* media (Gibco) supplemented with 10% heat-inactivated FBS (Sigma). S2 cells were transfected using the TransIT®-Insect Transfection Reagent, using 2 million cells per transfection and 1 µg of plasmid DNA, and collected after 48 hrs. For knockdown experiments, two rounds of electroporation (48 hrs apart) were performed using the Cell Line Nucleofector kit V (Amaxa Biosystems; program G-030), as described (Batki et al. 2019). Chromodomain constructs were co-transfected with 200 pmol of siRNA duplexes (oligo sequences are listed in **Table S4**) during the second nucleofection round or transfected using the TransIT®-Insect Transfection Reagent 24 hrs following the second nucleofection round. Cells were collected 5 days after the first nucleofection.

#### Western blot

Cell pellets and ovaries were lysed in RIPA buffer (Pierce) supplemented with protease inhibitors (Roche) and incubated for 30 min at 4 °C. After centrifugation at 21,000 g for 10 min at 4 °C, protein concentration was quantified using a Direct Detect Infrared Spectrometer (Merck). 20 µg total protein was separated on a NuPAGE 4–12% Bis-Tris denaturing gel (Thermo Fisher Scientific) and transferred to a nitrocellulose membrane using an iBLot2 dry transfer (Invitrogen). Primary antibody incubations were performed overnight at 4 °C and secondary antibodies were incubated for 1 hr at room temperature. All antibodies used are given in **Table S5**. Images were acquired using an Odyssey M Imaging system (LI-COR) and processed in Image Studio Lite (LI-COR).

### Data analysis

#### Processing of mRNA-seq and ribo-depleted RNA-seq

Adapters were removed using Trim Galore! (v0.6.4, --stringency 6) and with additional parameters “-a CTGTCTCTTATA --clip_R1 1 --clip_R2 1 --three_prime_clip_R1 1 --three_prime_clip_R2 1” if appropriate. The resulting reads were mapped to *dm6* using STAR (v2.7.3a, --outMultimapperOrder Random –outSAMmultNmax 1 --outFilterMultimapNmax 1000 --winAnchorMultimapNmax 2000 --alignSJDBoverhandMin 1 --sjdbScore 3) and a genome index built using NCBI RefSeq. Gene expression was quantified using featureCounts (v1.5.3, -s 2 -O --largestOverlap -Q 50) with gene models from Ensembl (release 97) or annotations corresponding to each TE consensus sequences (RepBase, downloaded 2022-04-20).

#### Processing of sRNA-seq

The analyses included nos-driven *E(z), kipf, w* knockdown (2-3 replicates each), nos-driven *E(z)*-*w* and *E(z)*-*kipf* double knockdown (2 replicates each), pTOsk-driven *E(z), rhi*, or *w* knockdown (2 replicates each), and previously published (Baumgartner et al. 2022) Gal4-driven *kipf, rhi* or *w* knockdown (1 replicate each) and Kipf-KO, rhi-KO and *w1118* flies (1 replicate each). Adapters were removed using Trim Galore! (v0.6.4, --stringency 6 --length 18 --max_length 29 -q 0). For previously published sRNA-seq, we further used ‘-a’ to specify the adapter, and ‘--clip_R1’ and ‘--three_prime_clip_R1’ to remove non-standard adapters and random nucleotides at the read ends (see **Table S7** for details). The resulting reads were mapped to *dm6* using Bowtie (v2.3.1, -S -n 2 -M 1 --best --strata --nomaqround --chunkmbs 1024 --no-unal). Gene expression was quantified using featureCounts (v1.5.3, -s 1 [or -s 2] -O --largestOverlap -Q 40) with gene models from Ensembl (release 97) or annotations corresponding to each TE consensus sequences (RepBase, downloaded 2022-04-20).

#### Differential gene expression and piRNA abundance analysis

RNA-seq differential expression analyses were performed using the DESeq2 package from R/Bioconductor. For gene expression, we first applied the DESeq2 function with default parameters followed by fold change shrinkage using the ‘ashr’ method. To analyse TE expression, we performed a similar analysis but used the size factors previously derived based on gene expression, while dispersion and fold change shrinkage was estimated based on both gene and TE expression to increase robustness in the estimates. RPKM values were calculated following the robust median implementation in DESeq2’s ‘fpm’ function. To consider a gene or TE to be differentially expressed, we required a 4-fold change and padj<0.05 unless otherwise specified.

sRNA-seq was analysed in a similar way, except that the analysis was restricted to reads of length 23-30 nt mapping sense or antisense, respectively, to the TE consensus sequences. Size factors were estimated separately using the estimateSizeFactors function with siRNAs mapping antisense to annotated genes as input.

#### Processing of CUT&RUN data

CUT&RUN data from S2 cells and *D melanogaster* ovaries were generated in 1-4 replicates per condition as specified in **Table S7**. Sequencing adapters were removed using Trim Galore! (v0.6.4, --paired --stringency 6). The resulting reads were mapped to the *dm6* reference genome using Bowtie (v1.2.3, -S -y -M 1 --best --strata --fr --minins 10 --maxins 600 --chunkmbs 2000 --nomaqround), reporting at most one hit for each read and considering insert sizes between 10 and 600nt. PCR duplicates were removed using MarkDuplicates from Picard tools (v2.21.2). All the following analyses used the deduplicated data, except for the genome browser visualization.

#### Processing of ChIP-seq data

ChIP-seq reads were aligned to the dm6 genome using Bowtie2 (v2.4.2), with the alignment process set to report at most one hit for each read. In case of alignments with the same MAPQ score, the best alignment was randomly selected from among those equally scored alignments. Peaks were called using MACS2 (v2.2.7.1) to capture narrow (-q 0.05 -g dm) and broad peaks (-q 0.05 -g dm --broad --broad-cutoff 0.1). Only uniquely mapped reads were used for the peak calling. As a control we used Input for each condition.

#### Processing of publicly available ChIP-seq data

Some publicly available ChIP-seq libraries used for comparison to CUT&RUN were analysed slightly differently. Specifically, HP1a ChIP-seq data for from ovaries was downloaded from GEO (accession GSE140539) (Zenk et al. 2021). The libraries were paired-end 2x101 nt and performed in two replicates. Rhi ChIP-seq samples from control, Rhi, and Moon knockout ovaries and two ChIP-seq input samples were downloaded from GEO (accession GSE97719) (Andersen et al. 2017). The Rhi libraries were paired-end 2×50 nt and performed in one replicate per condition. The input samples were 2x100 nt and performed in two replicates. Rhi and Kipferl ChiP-seq samples from controls and Rhi and Kipf MTD-Gal4-mediated knockdown ovaries and corresponding input samples were downloaded from GEO (accession GSE202468) (Baumgartner et al. 2022). This data was had variable read length (50, 74, or 100 nt) and was processed as single-end 50 nt. An overview of all ChIP-seq samples is available in **Table S7**. Sequencing adapters were removed using Trim Galore! (v0.6.4 or v0.6.6, --paired --stringency 6), or alternatively ‘--hardtrim5 50’. The resulting reads were mapped to the *dm6* reference genome using either Bowtie (v1.2.3, -S -y -M

1 --best --strata --fr --maxins 500 --chunkmbs 2000 --nomaqround), reporting at most one hit for each read. PCR duplicates were removed using MarkDuplicates from Picard tools (v2.21.2). All the following analyses used the deduplicated data, except for the genome browser visualization.

#### Peak calling using CUT&RUN or publicly available ChIP-seq data

Peaks were called using MACS2 to capture either narrow (callpeak -f BAMPE -g dm -q 0.01) or broad (callpeak -f BAMPE -g dm -q 0.05 --broad --broad-cutoff 0.1) peaks. Only uniquely mapped reads were used for the peak calling. As controls we used the corresponding IgG libraries for CUT&RUN, or input libraries for ChIP-seq, except for when Rhi ChIP-seq were compared to CUT&RUN and we used the Rhi knockout ChIP-seq as control. For histone modifications with two or more CUT&RUN replicates and CDs with three replicates, we derived a consensus peak set by first merging all peaks and then excluding peaks that were not supported by at least two replicates. For remaining CUT&RUN samples, we used all MACS2 peaks. Area-proportional Venn diagrams were created in R using the eulerr (v7.0.2) package to provide the best fit. Some low count overlaps might not be displayed due to geometric constraints. All overlaps are listed in the corresponding source data files.

#### Binning analysis in *Drosophila melanogaster*

The binning analysis was done using either 1 kb or 10 kb bins. For 1 kb bins, the genome was divided into 144,916 non-overlapping 1 kb bins. To avoid regions with low mappability, we excluded bins with less than 20% mappability (fewer than 200 mappable positions) resulting in 125,519 mappable 1 kb bins. For 10 kb bins, the genome was divided into 29,918 bins using a 10 kb sliding window that moves 5 kb at a time. Bins at the chromosome ends with size less than 5 kb were removed. To avoid regions with low mappability, we excluded bins with 20% mappability (fewer than 2,000 mappable positions) resulting in 25,865 mappable 10 kb bins. Bins derived from mitochondria were further removed, producing a final set of 125,499 bins of size 1 kb and 25,862 of size 10 kb for CUT&RUN. When integrating data from multiple assays, selection of mappable bins was done with respect for the CUT&RUN libraries, and the same bins were used for RNA-seq and sRNA-seq analyses. For binning analysis including only sRNA-seq or RNA-seq, we re-calculated mappability using bowtie alignment with either 100 nt or 26 nt read lengths. CUT&RUN and publicly available ChIP-seq signal was quantified per 50 nt window using the bamCompare module from deepTools (v3.3.2, --binSize 50 --ignoreForNormalization chrM -p 4 --scaleFactorsMethod SES --extendReads --centerReads --exactScaling --minMapping Quality 255 -of bedgraph --operation subtract --pseudocount 0) using pooled IgG CUT&RUN samples as background, except for Rhi ChIP-seq where we used the *rhi* knockout ChIP-seq as background. Next, values below zero were set to zero, before the average normalized signal per window was calculated using bedtools map. RNA-seq and sRNA-seq signal was quantified in a stand-specific manner by converting uniquely mapped reads to BED format and counting the number of reads from each strand falling into a bin with at least half of their length (bedtools intersect, -c -F 0.5).

#### Genome browser visualization

For CUT&RUN, RNA-seq and sRNA-seq, we first counted the number of all and uniquely mapped reads, respectively, using samtools, per strand if applicable. Uniquely mapped reads were converted into bigWig files using the deepTools bamCoverage module (v3.3.2, --binSize 1 –ignoreForNormalization chrM --normalizeUsing CPM --exactScaling --scaleFactor *s* --skipNonCoveredRegions --minMappingQuality 255), where the scale factor, *s*, was set to the number of uniquely mapped reads divided by all mapped reads. Additionally, we used ‘--extendReads --centerReads’ for CUT&RUN samples to centre the reads at fragments midpoints, ‘--filterRNAstrand’ for RNA-seq and sRNA-seq to separate the two strands, and ‘--minFragmentLength 23 --maxFragmentLength 30’ to select piRNAs for sRNA-seq. Uniquely mapped reads were converted into bigwig files using deepTools bamCoverage function (v3.5.0 - -binSize 1 --ignoreForNormalisation ChrM --normalizeUzing CPM --extendReads --centerReads --skipNonCoveredRegions).

#### Assessing classification performance using AUC

Classification performance for individual histone marks or combinations was assessed using a receiver operating characteristic (ROC) curve displaying true positive rate against false positive rate. Positive instances were defined as bins overlapping the cluster(s) of interest, with the remaining bins considered negative. The cumulative distribution was calculated in R using the ‘cumsum’ function with area under the curve calculated using the trapezoidal method.

#### Assessing Kipf-dependency of Rhi peaks

Rhi ChIP-seq peaks were derived using MACS2 as described above. To focus on the most reliable binding sites, we merged peaks with at least 3-fold enrichment over background in individual control knockdown replicates into a high-confidence peak set. Any peak that was located to an unplaced contig or that was also present in *rhi*-KD was excluded from the analysis. Finally, the remaining peaks were subdivided into Kipf-independent (n=463) or Kipf-dependent (n=1166) peaks, depending on whether they overlapped a Rhi peak in Kipf-KD or not.

#### Coverage plots at Rhi peaks

To visualize read coverage over Rhi peaks (**Fig. 3g**) we used the deepTools computeMatrix module (v3.3.2, reference-point --bin-size 50 -b 5000 -a 5000 --missingDataAsZero --reference-point center). The signal in the peak region was shown as difference between the knock-down and the control conditions.

#### Contribution of E(z) and Kipf, respectively, to piRNA production

Analysis was performed per 1 kb bin across the genome. First, we identified 632 bins where piRNA production depends on Rhi (>2-fold reduction in cpm, across pTOsk-Gal4- or MTD-Gal4-driven knockdown, and *rhi* knockout; 1-2 replicates each). Next, we calculated the change in piRNA abundance in E(z)- or Kipf-depleted ovaries (nos-Gal4- or pTOsk-Gal4-driven knockdown of *E(z)*, 2 replicates each; MTD-Gal4-driven knockdown of *kipf* or *kipf* knockout, 1 replicate each). Bins with >4-fold reduction in ovaries depleted for Kipf, E(z), or both were considered to be Kipf-dependent, E(z)-dependent or dependent on both.

#### Correlation heatmaps

Heatmaps were produced using the R package pheatmap (v1.0.12) with Euclidean distance.

#### Analysis of CUT&RUN in *E(z)* knockdown samples

To study the genome-wide effect of *E(z)* or *kipf* knockdown in S2 cells, we represented the genome as 117,300 1 kb bins and divided it into six equally sized groups based on the change in H3K27me3 signal upon E(z) loss. Four groups represented variable levels of H3K27me3 loss, one group represented no change, and one group displayed a relative gain in H3K27me3 signal. Next, across each group, we calculated the change in 2xPc, 2xCBX1, 2xRhi, and Pc-CBX1 binding affinity. Although 2xPc was strongly responsive to H3K27me3 loss, the other chromatin binders were largely unaffected. Hence, we next restricted the analysis to 46,583 bins across the six groups with H3K9me3 (defined as CD^2xCBX1^ above 90th percentile of euchromatic regions).

#### Binding affinity analysis using CUT&RUN data

Differential binding affinity analysis was performed using DiffBind (v3.8.4) applied on the MACS2 narrow or broad peaks from *siRen* and *siEz* samples, using the corresponding IgG libraries as controls and without specifying any blacklist regions. For the analysis, we used the dba, dba.count, dba.normalize, dba.contrast, and dba.analyze modules with default options. This will create a consensus set of peaks present in at least two samples, and then re-quantify the signal intensity at each consensus peak, and re-center and re-size the peaks to a 401nt region around their maxima. This resulted in 32,042 narrow consensus or 11,053 broad consensus peaks. The contrasts were specified as *siE(z)* against *siRen* for each target (H3K9me3, H3K27me3, CD^2xPc^, CD^2xCBX1^, CD^Pc-CBX1^, CD^2xRhi^).

#### Euchromatin and heterochromatin coordinates

We used the following coordinates from Fabry and colleagues (Fabry et al. 2021) to define euchromatin (chr2R:6460000-25286936, chr2L:1-22160000, chr3L :1-23030000, chr3R:4200000-32079331, chrX:250000-21500000) and heterochromatin (chr2R:1-6460000, chr2L:22160000-23513712, chr3L:23030000-28110227, chr3R:1-4200000) in *D melanogaster*.

#### Reference genomes

We used the *dm6* assembly for *D. melanogaster* downloaded from the UCSC genome browser and the following assemblies downloaded from NCBI: GCF_003285975 for *D. annanassae*, GCF_003286155 for *D. erecta*, GCF_016746395 for *D. simulans*, and GCF_016746365 for *D. yakuba*.

#### Genome-wide mappability

To estimate genome mappability, we divided the genome into all possible *n*-mers, where *n* is either 26 (for sRNA) or 100 (for CUT&RUN). Those sequences were then mapped back onto the genome and mappability for each position was estimated as the number of reads overlapping a position, divided by *n*.

#### Analysis of CUT&RUN in five *Drosophila* species

CUT&RUN was performed in *D. annanassae, D. melanogaster, D. erecta, D. simulans*, and *D. yakuba* for H3K27me3 and H3K9me3 with two replicates per condition, followed by 2×50 nt paired-end sequencing. Sequencing adapters were removed using Trim Galore! (v0.6.4, --paired --stringency 6 -a GATCGGAAGAGCACACGTCTGAACTCCAGTCAC). The resulting reads were mapped to their respective reference genome using Bowtie (v1.2.3, -S -y -M 1 --best --strata --fr --minins 10 --maxins 600 --chunkmbs 2000 --nomaqround), reporting at most one hit for each read and considering insert sizes between 10 and 600 nt. One of the H3K9me3 replicates for *D. yakuba* displayed very low complexity (1.3 million uniquely mapped reads with estimated library size 1.7 million fragments) and was therefore excluded from the analysis.

#### Analysis of sRNA-seq in five *Drosophila* species

Small RNA-seq was performed in *D. annanassae, D. melanogaster, D. erecta, D. simulans*, and *D. yakuba* with three replicates per species. The libraries were sequenced as 2×50 nt paired-end sequencing, but only the first read was used for the analysis. Trim Galore! (v0.6.4) was used to remove an abundant rRNA (--stringency 30 -a TGCTTGGACTACATATGGTTGAGGGTTGTA --length 18 -q 0) and sequencing adapters (--stringency 5 -a TGGAATTCTCGG --length 18 --max_length 35 -q 0). Next, Bowtie was used to exclude reads mapping to known *Drosophila* viruses (v.2.3.1, -S -M 1 --best --strata --nomaqround --chunkmbs 1024) using the ‘--max and --un’ options to extract unmapped reads. These reads were mapped to their respective reference genome (-S -M 1 --best --strata --nomaqround --chunkmbs 1024), reporting at most one hit for each read.

#### Binning analysis in five *Drosophila* species

For the binning analysis, the genome was divided into either 1 kb non-overlapping bins or 10 kb bins with 5 kb overlap. To estimate the mappability within each bin, we divided the genome into all possible 100-mers and mapped those sequences back onto the genome. Mappability was calculated as the number of 100-mers mapping uniquely within each bin divided by the bin size. Bins with size <5 kb (for 10 kb bins) or with <20% mappability were excluded, leaving between 121,081 and 178,773 1 kb or between 24,687 and 38,112 10 kb bins per species. We next quantified the number of uniquely mapped CUT&RUN or sRNA-seq reads per library mapping ≥50% within each bin, considering the forward and reverse strand separately for the sRNA-seq data. To correct for differences in sequencing depth, mappability, and bin size, the resulting bin counts were normalized by their mappability and converted to fpkm values. Bins from mitochondria were excluded from all subsequent analyses.

#### Scatterplots and violin plots in five *Drosophila* species

The normalized rpkm values were capped at 300 (or 200 for 10 kb bins) and log_10_-transformed using a pseudo-count of 1. To focus on bins with piRNAs derived from both strands, we represented the sRNA-seq data as minimum signal on the forward and reverse strands. For H3K9me3 and H3K27me3 we used the mean of two replicates and for sRNA-seq we used the mean of three replicates. For the violin plots, we grouped the bins based on their piRNA level and displayed the piRNA, H3K27me3 or H3K9me3 level within each group. Briefly, we first constructed one group for bins with no piRNAs. Next, we sorted all remaining bins by piRNA level and extracted 15 equidistant breakpoints, including the lowest and highest piRNA level. Each pair of consecutive non-identical breakpoints were used to construct a piRNA level interval, resulting in between 11 or 14 additional groups per species (1 kb) or between 14 and 15 additional groups (10 kb). All scatterplots and violin plots were constructed in R (v3.6.2) using base graphics, imageScale from sinkr (v0.6), gridExtra, and ggplot2.

#### Structure modelling of Rhi dimer using AlphaFold multimer

AlphaFold v2.3.2 Colab was used to generate a prediction model (Jumper et al. 2021). The full-length Rhi chromodomain protein (amino acids 20-90) reported in the crystal structure of **4U68** was used as input twice to generate the multimer (Yu et al. 2015). The structure of the Rhi chromodomain dimer was found to resemble the original structure of **4U68** well (Yu et al. 2015).

#### Structure modelling of histone 3 peptide using Zdock

Zdock was used to evaluate potential binding modes of the peptide to the dimer. To generate the structure of the input histone 3 peptide, the existing crystal structure of the K9me3-and K27me3-containing peptides in **4U68** and **1PFB** were used (Min et al. 2003; Yu et al. 2015). Within the structure of **4U68**, information for the structure of KQTARK(Me3)S, of the K9me3 section of the histone peptide was available. The structure of LATKAAR(Me3)SAP, of the K27me3 section of the histone peptide was available via **1PFB**. A structure alignment was done using **4U68** and **1PFB** via PyMOL, and the connecting chain of ‘TGGKAPRKQ’ was added using Modeller tool in UCSF Chimera 1.16 (Fiser and Sali 2003; Pettersen et al. 2004). This peptide was then docked using Zdock server, with ZDOCK 3.0.2 (Pierce et al. 2014) to yield a predictive model of the binding motif.

#### Molecular dynamics simulations

The structural prediction from AlphaFold multimer was used as a basis for molecular dynamic (MD) simulations. MD simulations in explicit water were performed using the graphics processing unit accelerated code (PMEMD) of the Amber 16 and AmberTools 20 packages (Case et al. 2016; Case et al. 2020). Protonation states were calculated using the PDB2PQR server. For the protein scaffold, an evolved version of the Stony Brook modification of the Amber 99 force field (ff14SB) (Maier et al. 2015) parameters were applied (see below for additional non-standard parameters). TIP3P parameters were assigned to water molecules (Jorgensen et al. 1983). To incorporate the non-canonical amino acid K(Me3), into the AlphaFold model, the structure of the K(Me3) residue was used from **4U68** and the partial charges of the K(Me3) residue were set to fit the electrostatic potential generated at the HF/6-31G(d) level by the restrained electrostatic potential model (Bayly et al. 1993). The charges were calculated according to the Merz–Singh–Kollman scheme using Gaussian (Besler et al. 1990). Each protein complex was immersed in a pre-equilibrated cubic box with a 12 Å buffer of TIP3P water molecules using the Leap module. The systems were neutralized by the addition of explicit counterions (Na^+^ or Cl^−^). Long-range electrostatic effects were modelled using the particle mesh Ewald method with periodic boundary conditions (Darden et al. 1993). An 8 Å cut-off was applied to Lennard-Jones and electrostatic interactions.

MD simulations were performed according to the following steps: (1) Minimization was performed with a maximum cycle of 5000 and with the steepest descent algorithm for the first 2500 cycles, with a periodic boundary for constant volume (canonical ensemble, NVT) and without the SHAKE (an algorithm for constrained molecular dynamics) algorithm activated. Positional restraints of 2 kcalmol^−1^Å^−2^ were applied on heavy atoms of the protein backbone and heavy atoms of the ligand. (2) A 1 ns heating process was performed with a periodic boundary for constant volume (NVT) with the SHAKE algorithm turned on such that the angle between the hydrogen atoms was kept fixed. Temperature increased from 0 K to 300 K in a time period of 1 ns with heat bath coupling with time constant of 2 ps. Positional restraints of 2 kcalmol^−1^Å^−2^ were applied on heavy atoms of the protein backbone. (3) A 2 ns equilibrium process was performed with a periodic boundary for constant volume (NVT) with the SHAKE algorithm turned on such that the angle between the hydrogen atoms was kept fixed. An Andersen-like temperature coupling scheme is used to maintain the temperature at 300 K. (4) A 2 ns equilibrium process was performed with a periodic boundary for constant pressure (isothermal-isobaric ensemble, NPT) and with a constant temperature of 300 K, maintained using Langevin dynamics with the collision frequency of 5 ps^−1^. (5) A 100 ns equilibrium process was performed with a periodic boundary for constant pressure (NPT) and with a constant temperature of 300 K. (6) A 1000 ns production was performed with a periodic boundary for constant pressure (NPT) and with a constant temperature of 300 K. A representative frame was obtained from 1 µS MD simulations using Chimera’s cluster analysis tool (Kelley et al. 1996).

## Data availability

Sequencing data generated in this study has been deposited to the GEO under accession (GSExxxxxx).

## ACKNOWLEDGEMENTS

We thank members of the Brasset and Hannon groups for fruitful discussions. We thank Silke Jensen for her help with preliminary RNA-seq analysis. We thank Lisa Baumgartner and Julius Brennecke for sharing unpublished data and scientific discussion. We thank Marie Bao from the Life Science Editors for feedback and comments on the manuscript. We thank the Scientific Computing, Genomics, RICS and microscopy core facilities at CRUK Cambridge Institute and the CLIC facility (Clermont Imagerie Confocale) at iGReD for support. GJH is a Royal Society Wolfson Research Professor (RSRP\R\200001). Work in the Hannon lab was funded in whole, or in part, by Cancer Research UK (G101107) and the Wellcome Trust (110161/Z/15/Z and 226627/Z/22/Z). PC and MJ were supported by Cancer Research UK grants C9685/A26398, C9685/A27415 and C9545/A29580. Work in the Brasset lab was funded by the Agence Nationale pour la Recherche (CHApiTRE) ANR-20-CE12-0005, BiopiC ANR-21-CE12-0022, and EpiTET 699 ANR-17-CE12-0030-03), the La Fondation ARC pour la recherche sur le cancer (PJA20171206129). This research was financed also by the French government IDEX-ISITE initiative 16-IDEX-0001 (CAP 20-25).

## AUTHOR CONTRIBUTIONS

A.A. and E.K. performed all experiments except those stated below. S.B. performed all computational analyses except ChIP-seq analysis, and initial RNA-seq and small RNA-seq analyses which were performed by Y.R. and S.M.-M., and the MD simulations/modelling which was performed by M.J. under the supervision of P.C.. E.L.E. performed knockdown experiments using pTOsk-Gal4 and generated RNA-seq libraries, J.v.L. generated small RNA-seq libraries. A.S. and Y.G. performed CUT&RUN from CD constructs in S2 cells, and Y.G. performed Western blot analysis. N.G. generated libraries for RNA-seq and prepared probes for RNA-FISH. The project was conceived by A.A., E.K., B.C.N., E.B. and G.J.H., and supervised by B.C.N., E.B. and G.J.H. The data was analysed and interpret by A.A., E.K., S.B., B.C.N., E.B. and G.J.H.. The paper was written by A.A., E.K., S.B., B.C.N., E.B. and G.J.H. with input from all authors.

