## Supplemental Figures and Notes for "A dual histone code specifies the binding of heterochromatin protein Rhino to a subset of piRNA source loci"

**Supplementary Figures**

**Extended Data Fig. 1 (related to Fig. 1):** Summary of screen results.
**Extended Data Fig. 2 (related to Fig. 1):** *E(z)*-depleted ovaries display piRNA loss and TE de-repression.
**Extended Data Fig. 3 (related to Fig. 2):** Rhi colocalises with H3K9me3 and H3K27me3 in germ cells/Presence of H3K9me3 and H3K27me3 is predictive on dual-strand cluster loci.
**Extended Data Fig. 4 (related to Fig. 3):** Rhi binding at Kipf-independent loci depends on E(z)
**Extended Data Fig. 5 (related to Fig. 3):** Rhi binding upon *E(z)-kipf* double knockdown
**Extended Data Fig. 6 (related to Fig. 4):** Rhi chromodomains associate with regions co-occupied by H3K9me3 and H3K27me3
**Extended Data Fig. 7 (related to Fig. 4):** $CD^{2xRhi}$ binding in S2 cells depends on E(z)
**Extended Data Fig. 8 (related to Fig. 5):** Dual-strand piRNA producing loci are associated with both H3K9me3 and H3K27me3 in Drosophilids

**Supplementary Tables**

**Table S1:** Table showing small RNA-seq differential expression analysis data used for plotting.
**Table S2:** Table showing RNA-seq differential expression analysis data used for plotting.
**Table S3:** List of fly lines and species used in this study.
**Table S4:** List of oligonucleotides used in this study.
**Table S5:** List of antibodies used in this study.
**Table S6:** Sequences of chromodomain constructs used in this study.
**Table S7:** List of all sequencing libraries analysed in this study and their alignment metrics.

**Supplementary Notes**

**Supplementary Note 1 (related to Extended Data Fig. 1):** PRC2 components recovered in screen
**Supplementary Note 2 (related to Fig. 2):** Antibody cross-reactivity

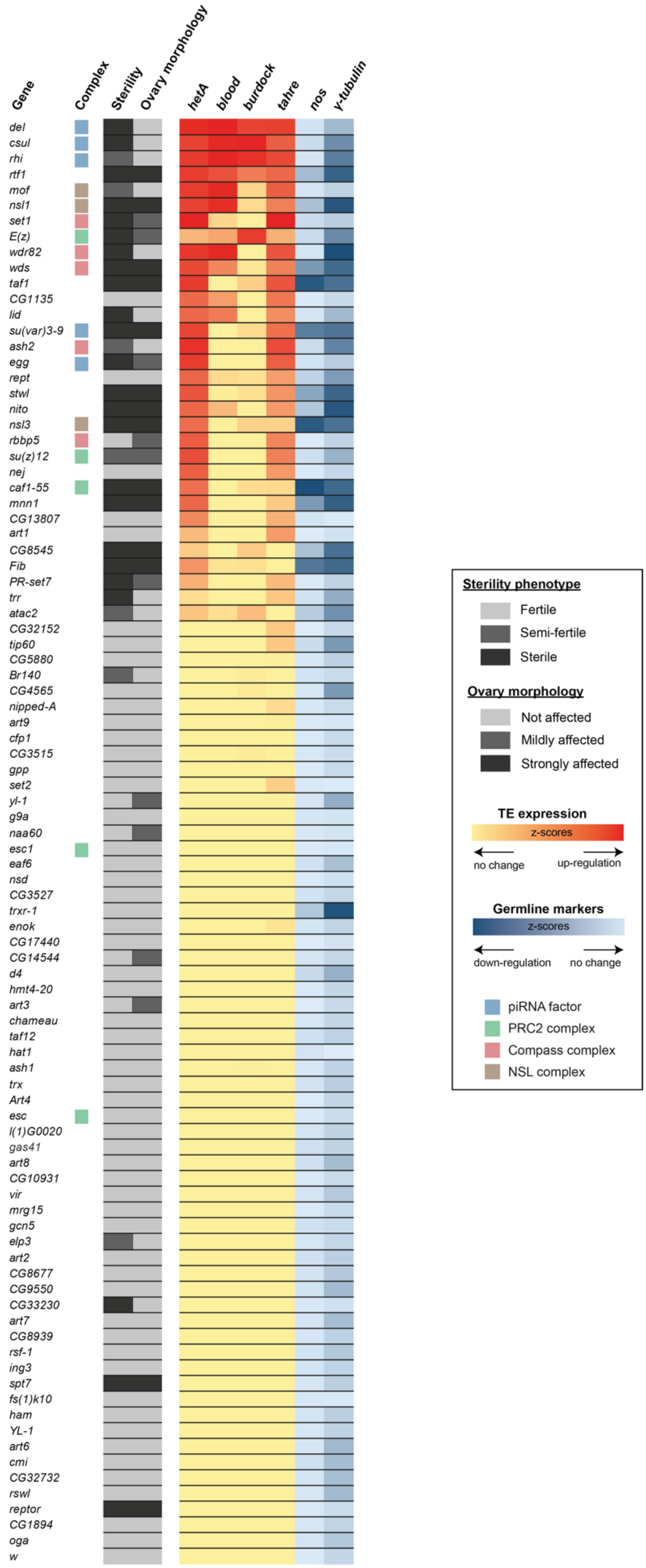


**Extended Data Fig. 1 (related to Fig. 1): Summary of screen results.**
Heatmaps summarising gene names, fertility and ovary morphology phenotypes, as well as TE (*hetA*, *blood*, *burdock*, *tahre*) and germline marker (*nos*, *yTub*) expression, measured by RT-qPCR, upon indicated germline knockdowns (displayed as z-scores). Known piRNA factors and proteins that are part of the COMPASS, PRC2 and NSL complexes are highlighted as indicated in the legend.

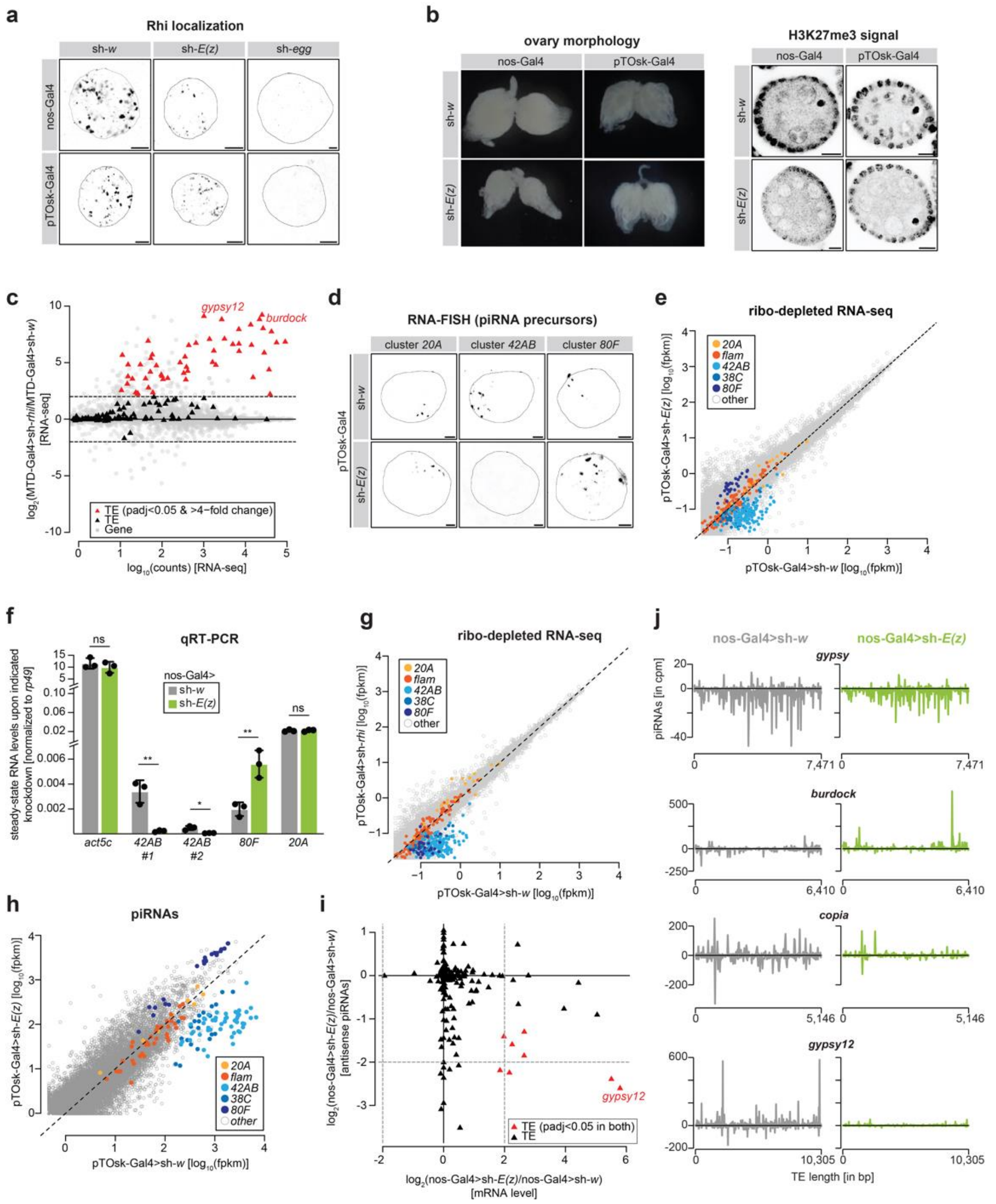


**Extended Data Fig. 2 (related to Fig. 1): *E(z)*-depleted ovaries display piRNA loss and TE de-repression.** **a,** Immunofluorescence staining showing Rhi expression and localisation in nurse cell nuclei of stage 4 egg chambers upon the indicated shRNA and Gal4 driver combination (scale bar: 5 µm). **b,** Ovary morphology image (left) and confocal images of egg champers stained with anti-H3K27me3 (right) upon the indicated knockdowns and drivers (scale bars: 10µm). **c**, MA plot showing counts per gene (grey) and TEs (black) in polyA-selected RNA-seq libraries from ovaries of MTD-Gal4 mediated germline knockdown of *rhi* versus control. TEs with padj<0.05 and >4x fold change are shown in red. **d,** Confocal images showing RNA-FISH signal for transcripts derived from piRNA clusters *42AB*, *80F* and *20A* in control and E(z) depleted ovaries using the pTOsk-Gal4 driver (scale bar: 5 µm). **e,** Scatter plot depicting normalized ribo-depleted RNA levels (fpkm) of uniquely mapping reads in 1 kb bins in ovaries with pTOsk-Gal4 driven *E(z)* knockdown versus control (average of four replicate experiments each). **f,** Bar graphs showing levels of the indicated piRNA precursor transcripts measure using RT-qPCR upon the indicated germline knockdown. Data are presented as the mean (n=3) with error bars representing standard deviation. Statistical significance was determined using the unpaired t-test. ** corresponds to P-value < 0.01, * corresponds to P-value < 0.05, and ns corresponds to p>0.05. **g**, same as e but showing pTOsk-Gal4 driven *rhi* knockdown versus control (average of four replicate experiments each). **h**, Scatter plot depicting normalized piRNA levels of uniquely mapping piRNAs in 1 kb bins in ovaries with pTOsk-Gal4 driven *E(z)* knockdown versus control (two replicates each). **i**, Scatter plot showing TE transcript fold-change against antisense piRNA fold-change in nos-Gal4 driven *E(z)* knockdown versus control. Filled triangles indicate TEs that change on both mRNA and piRNA level (DESeq2, padj<0.05). **j,** Profile plots of piRNA reads over selected TE consensus sequences (*burdock*, *copia*, *gypsy12* and *gypsy*). piRNA counts (normalized to miRNAs) are displayed for indicated genotypes.

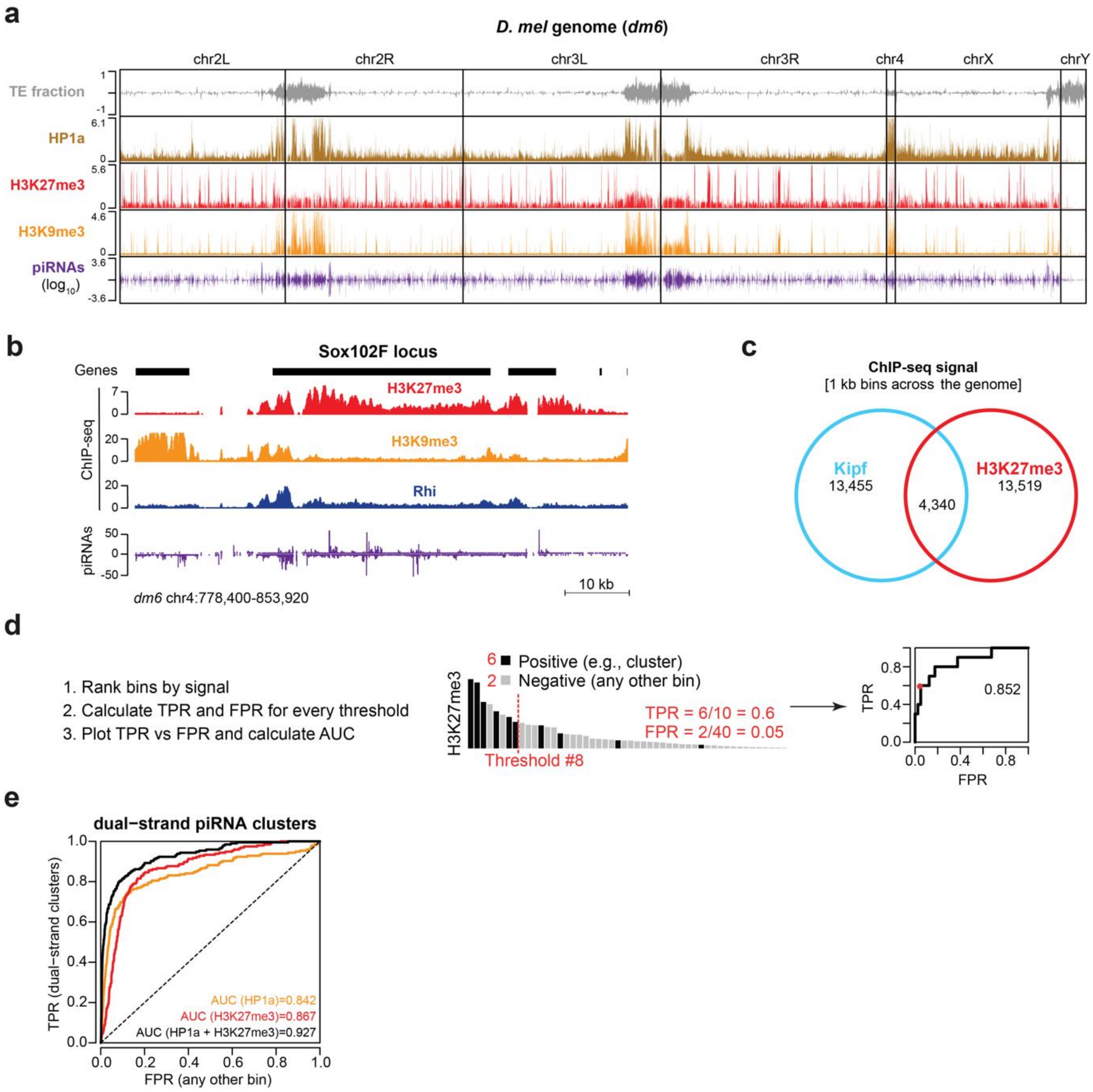


**Extended Data Fig. 3 (related to Fig. 2): Rhi colocalises with H3K9me3 and H3K27me3 in germ cells.**
**a,** Genome-wide view of TE content across *D. melanogaster* chromosomes with TE fraction (grey), HP1a (beige) ChIP-seq (Zenk et al. 2021), H3K27me3 (red) and H3K9me3 (yellow) CUT&RUN and piRNA levels (purple) in wildtype ovaries. CUT&RUN was normalized to IgG signal per 50 nt (see **Methods** for details). **b,** UCSC genome browser tracks of ChIP-seq signal (coverage per million reads) of Rhi (blue), H3K27me3 (red), and H3K9me3 (yellow), and uniquely mapping piRNAs (purple, normalized to miRNA reads) at the *sox102F* locus in nos-Gal4>sh-*w* ovaries. **c,** Venn diagram showing overlap between Kipf and H3K27me3 ChIP-seq signal across genomic 1 kb bins. Signal was considered to be present if the bin overlapped a MACS2 broad peak (q<0.05, broad-cutoff<0.1) across two pooled biological replicates. **d,** Schematic showing how predictive performance was assessed in a threshold-independent way using an area under ROC curve metric. First, all genomic 1 kb bins were ranked according to signal strength (e.g., highest to lowest H3K27me3). Second, for every possible rank threshold, true positive rate (TPR) is calculated as number of bins of interest (e.g., located within a dual-strand piRNA cluster) above the threshold divided by the total number of bins of interest. Similarly, false positive rate (FPR) is calculated as the number of other bins (e.g., located outside of a dual-strand piRNA cluster) above the threshold, divided by the total number of other bins. Third, TPR is plotted against FPR and the area under curve (AUC) is calculated. A higher metric indicates more predictive power and AUC at 0.5 indicate performance expected by random guessing. **e,** Line graphs comparing the ability to identify dual-strand piRNA clusters based on the strength of individual features (H3K27me3 and HP1a) or combinations.

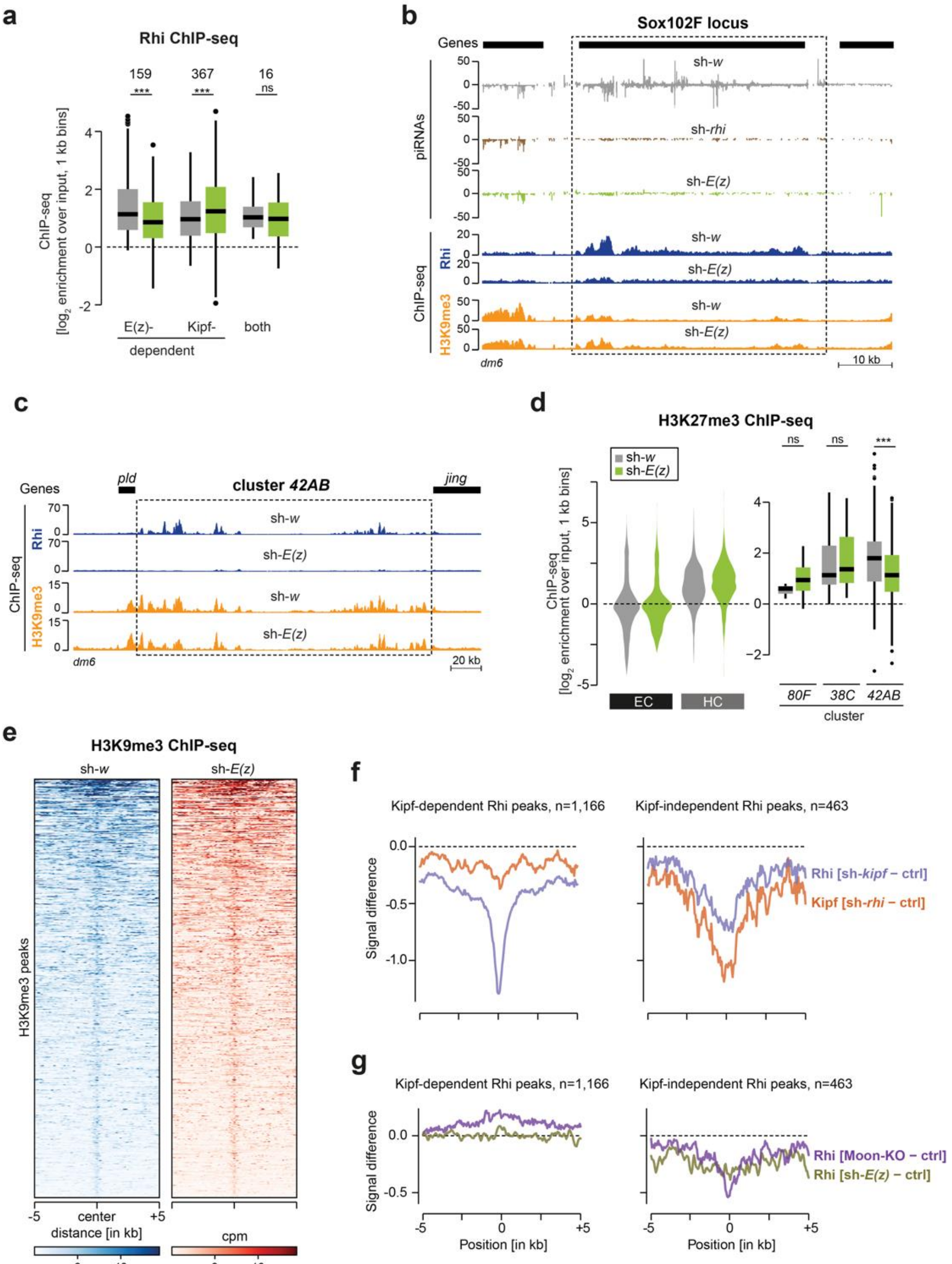


**Extended Data Fig. 4** (**related to Fig. 3**)**: Rhi binding at Kipf-independent loci depends on E**(**z**).
**a**, Box plots showing Rhi ChIP-seq enrichment per E(z)/Kipf-dependency category. *** corresponds to P<0.001, ns corresponds to p>0.05 based on Wilcoxon signed-rank test. **b,** UCSC genome browser tracks of the *sox102F* locus displaying profiles of small-RNAs uniquely mapping in control, *E(z)*-depleted and *Rhi*-depleted ovaries and ChIP-seq signal for Rhi and H3K9me3 in control and *E(z)*-depleted ovaries. **c**, UCSC genome browser tracks of ChIP-seq signal for Rhi and H3K9me3 in control and *E(z)*-depleted ovaries of the *42AB* dual-strand piRNA cluster. Dashed line indicates approximate piRNA cluster boundaries. **d,** Violin plots (left) and box plots (right) showing average $\log_2$-fold H3K27me3 enrichment by ChIP-seq from ovaries with nos-Gal4 driven *E(z)* knockdown versus control (average of two replicate experiments each) in heterochromatin (HC) and euchromatic chromosome arms (EC), quantified across 1 kb bins (excluding piRNA clusters). H3K27me3 occupancy at indicated piRNA clusters is shown as box plot quantification (n depicts 1 kb bins analysed for each piRNA cluster). *** corresponds to P<0.001, ns corresponds to p>0.05 based on Wilcoxon signed-rank test. Box plots show median (centre line), with interquartile range (box) and whiskers indicating at most 1.5x interquartile range). **e**, Heatmap of H3K9me3 ChIP-seq signal across wildtype H3K9me3 peaks (n= 4,785) in control and *E(z)* knockdown ovaries. **f**, Metaplot showing mean difference in ChIP-seq signal for Kipf (orange) in *rhi* knockdown, and Rhi (light blue) in *kipf* knockdown across Rhi peaks categorised as either Kipf-dependent or not (see **Methods**). **g**, Metaplot showing mean difference in ChIP-seq signal for Rhi in *moon* (purple) knockout or *E(z)* knockdown (yellowish green) across Rhi peaks categorised as either Kipf-dependent or not (see **Methods**).

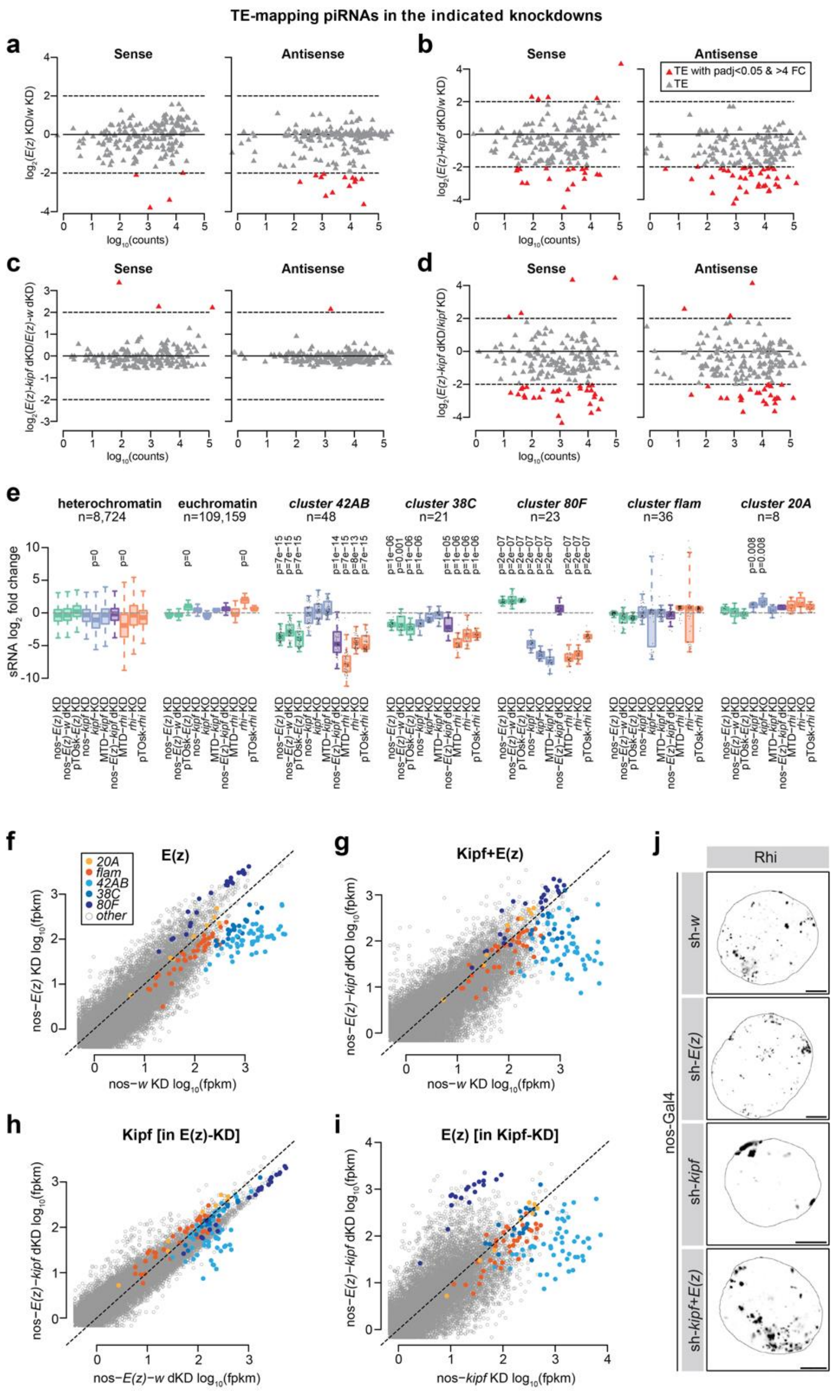


**Extended Data Fig. 5 (related to Fig. 3): Rhi binding upon *E(z)-kipf* double knockdown.**
**a-d,** MA plot showing counts per TE (black) in sRNA-seq libraries from ovaries of germline (**a**) *E(z)* KD versus control, (**b**) *E(z)-kipf* dKD versus control, (**c**) *E(z)-kipf* dKD versus *E(z)* KD, and (**d**) *E(z)-kipf* dKD versus *kipf* KD. Sense and antisense piRNAs are shown separately and TEs with padj<0.05 and >4x fold change are shown in red. **e,** Boxplots showing changes in piRNAs mapping uniquely to genomic 1kb bins within heterochromatin, euchromatin or indicated piRNA clusters (*n* refers to the number of 1kb bins for each category). Signal shown as a $\log_2$-ratio between the indicated KD and its respective control (average of 1-3 biological replicates). P-values were calculated using a two-sided paired Wilcoxon signed-rank test for >2-fold changes. **f,** Scatter plot depicting normalized uniquely mapping piRNA levels in 1 kb bins in ovaries with nos-Gal4 driven *E(z)* knockdowns versus control (average of two replicates each). **g,** Scatter plot depicting normalized uniquely mapping piRNA levels in 1 kb bins in ovaries with nos-Gal4 *E(z)-Kipf* dKD compared to nos-Gal4 driven *w* knockdown (average of two replicates each). **h,** Same as d but comparing *E(z)-kipf* dKD to nos-Gal4 driven *E(z)-w* dKD (average of two replicates each). **i,** Same as d but comparing *E(z)-kipf* dKD to nos-Gal4 driven *kipf* knockdown (average of two or three replicates each). **j,** Immunofluorescence staining showing Rhi expression and localisation in nurse cell nuclei upon the indicated shRNA and nos-Gal4 driver combination (scale bar: 5 μm).

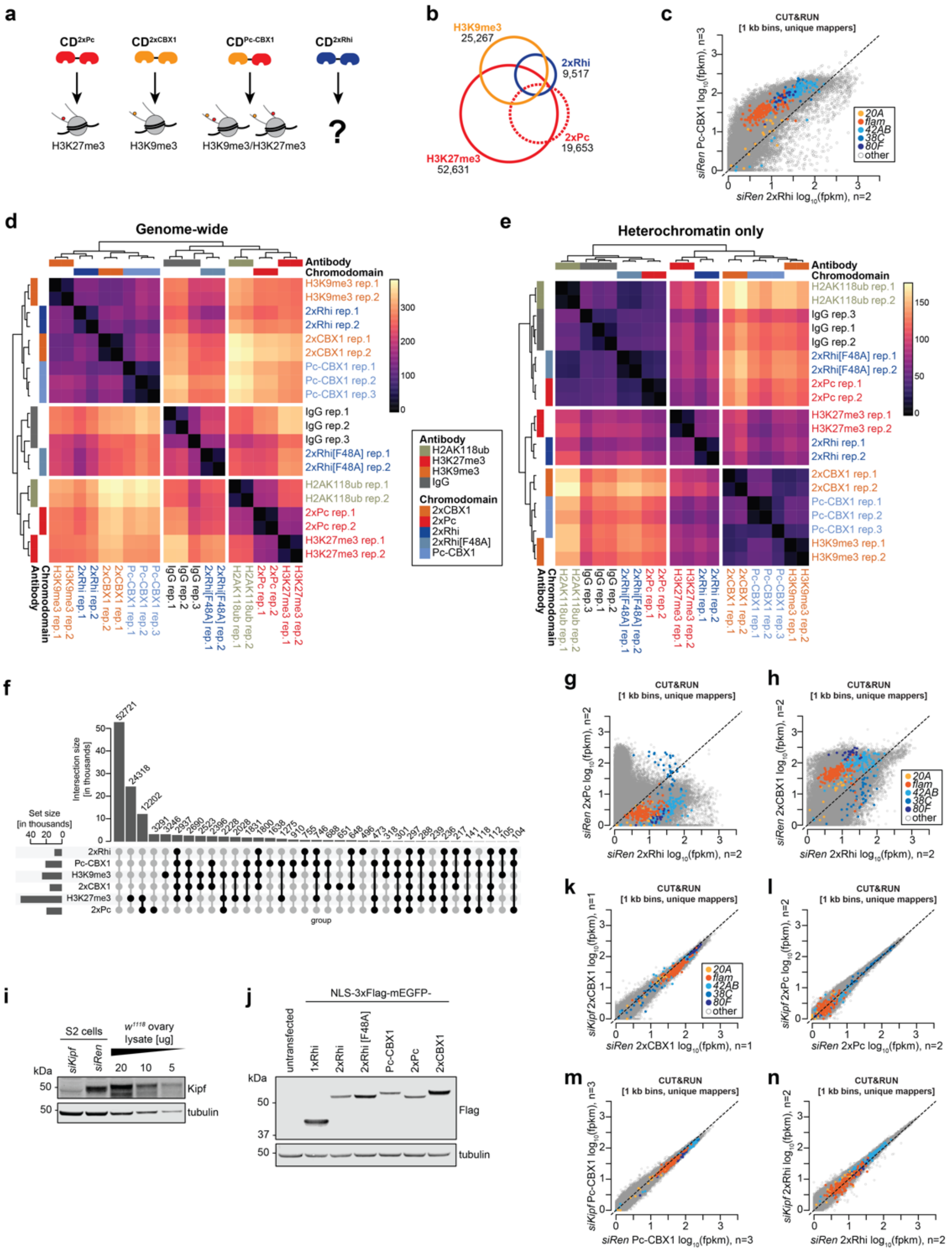


**Extended Data Fig. 6 (related to Fig. 4): Rhi chromodomains associate with regions co-occupied by H3K9me3 and H3K27me3.**

**a,** Schematic showing the experimental workflow and cartoon of the dual-CD constructs used in the assay. **b,** Venn diagram showing overlap between $CD^{2xPc}$ binding, H3K27me3, H3K9me3, and Rhi across 125,499 genomic 1 kb bins. Signal was considered to be present if the bin overlapped a MACS2 broad peak (q<0.05, broad-cutoff<0.1) present in at least two biological replicates. **c,** Scatter plot depicting CUT&RUN signal (fpkm) of uniquely mapping reads in 1 kb bins of $CD^{2xRhi}$ versus $CD^{Pc\text{-}CBX1}$ in S2 cells (average of 2-3 replicate experiments each). **d,** Heatmap and hierarchical clustering (Euclidian distance) of CUT&RUN signal detected for the indicated chromodomain constructs and histone modifications [$\log_{10}$(fpkm)]. **e**, Same as d, but across a subset of the genome classified as heterochromatin (10,180 out of 125,499 1 kb bins). **f,** UpSet plot showing all major intersections (>100 1 kb bins) across the six indicated CUT&RUN experiments. **g,** as in c but showing $CD^{2xRhi}$ versus $CD^{2xPc}$ (average of 2 replicate experiments each). **h,** as in c but showing $CD^{2xRhi}$ versus $CD^{2xCBX1}$ (average of 2 replicate experiments each). **i**, Western blot analyses showing Kipf expression in S2 cells (treated as indicated) or $w^{1118}$ ovaries. Tubulin is shown as loading control. **j**, Western blot analyses showing the levels of the indicated chromodomain constructs (Flag) in S2 cells that were transfected with the corresponding plasmids. Tubilin is shown as loading control. **k,** Scatter plot depicting CUT&RUN signal (fpkm) of uniquely mapping reads in 1 kb bins of $CD^{2xCBX1}$ upon control or *kipf* knockdown in S2 cells (1 replicate experiment).**l-**

**n,** as in k but depicting $CD^{2xPc}$, $CD^{Pc\text{-}CBX1}$, and $CD^{2xRhi}$, respectively (2-3 replicate experiments each, as indicated).

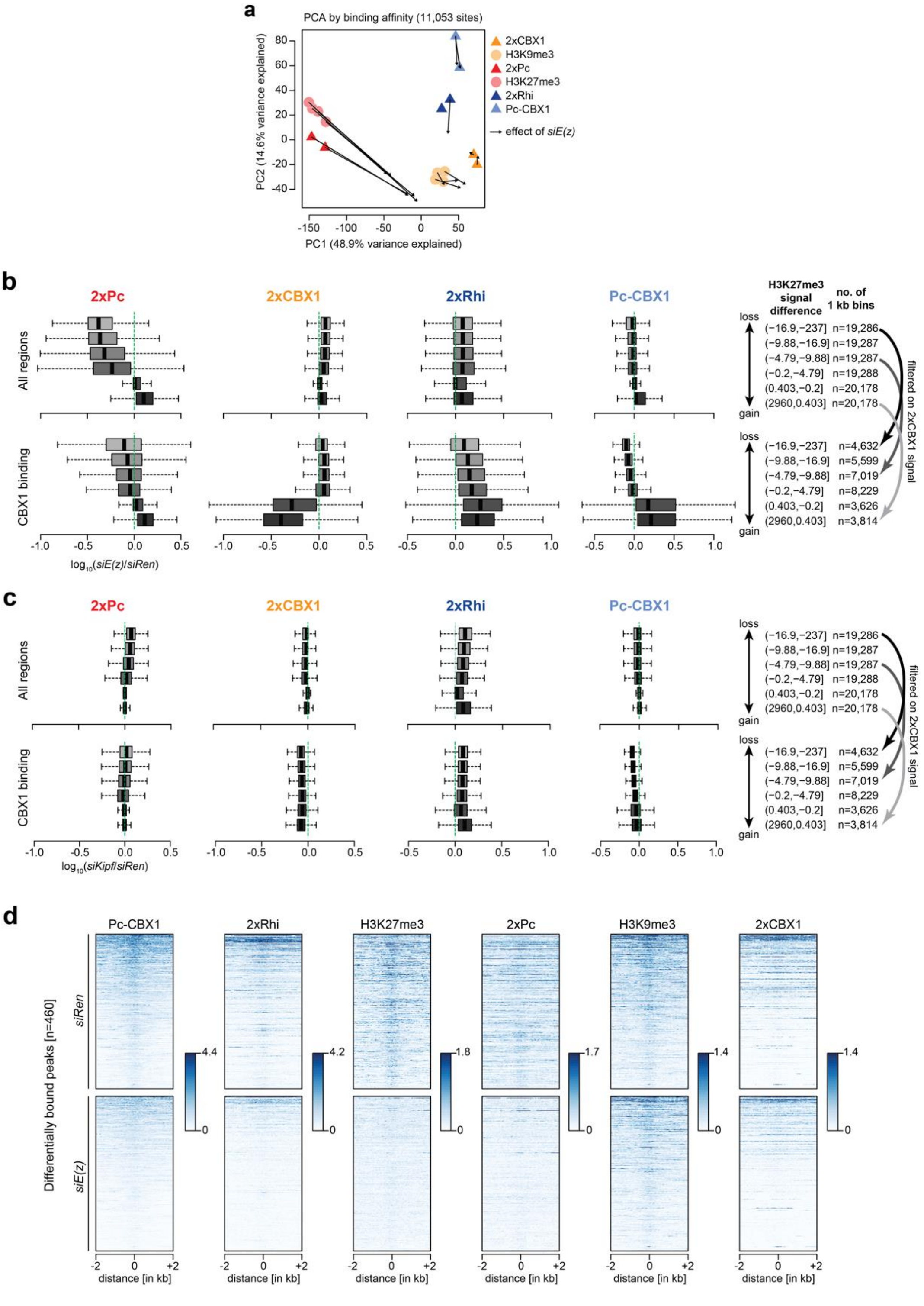


**Extended Data Fig. 7 (related to Fig. 4): CD$^{2xRhi}$ binding in S2 cells depends on E(z).**

**a,** Principal component analysis (PCA) plot of the binding affinities of the indicated chromodomain fusion proteins expressed in S2 cells in control condition (circle/triangle) and *E(z)* knockdown (arrowhead). One 2xRhi sample without arrowhead lacks a corresponding knockdown sample. **b,** Box plot showing loss of binding of the indicated chromodomain fusion proteins in S2 cell upon *E(z)* knockdown compared to control knockdown (*siRen*). Shown are $\log_{10}$ fold change per 1 kb bin either genome wide (top, n=117,300 bins with a non-zero change) or at CD$^{2xCBX1}$-bound regions (bottom, n=46,583 bins, CD$^{2xCBX1}$ above 90$^{th}$ percentile of euchromatic regions). The genome-wide bins were divided into six groups according to the severity of H3K27me3 loss (higher to lower, thresholds indicated to the right). **c**, Same as a, but for *kipf* knockdown (*siKipf*) compared to control knockdown (*siRen*). **d,** Heatmaps showing the indicated chromodomain fusion protein and histone modification CUT&RUN signal in the 4 kb surrounding CD$^{Pc\text{-}CBX1}$ differentially bound peaks in control (*siRen*) and *E(z)* knockdowns.

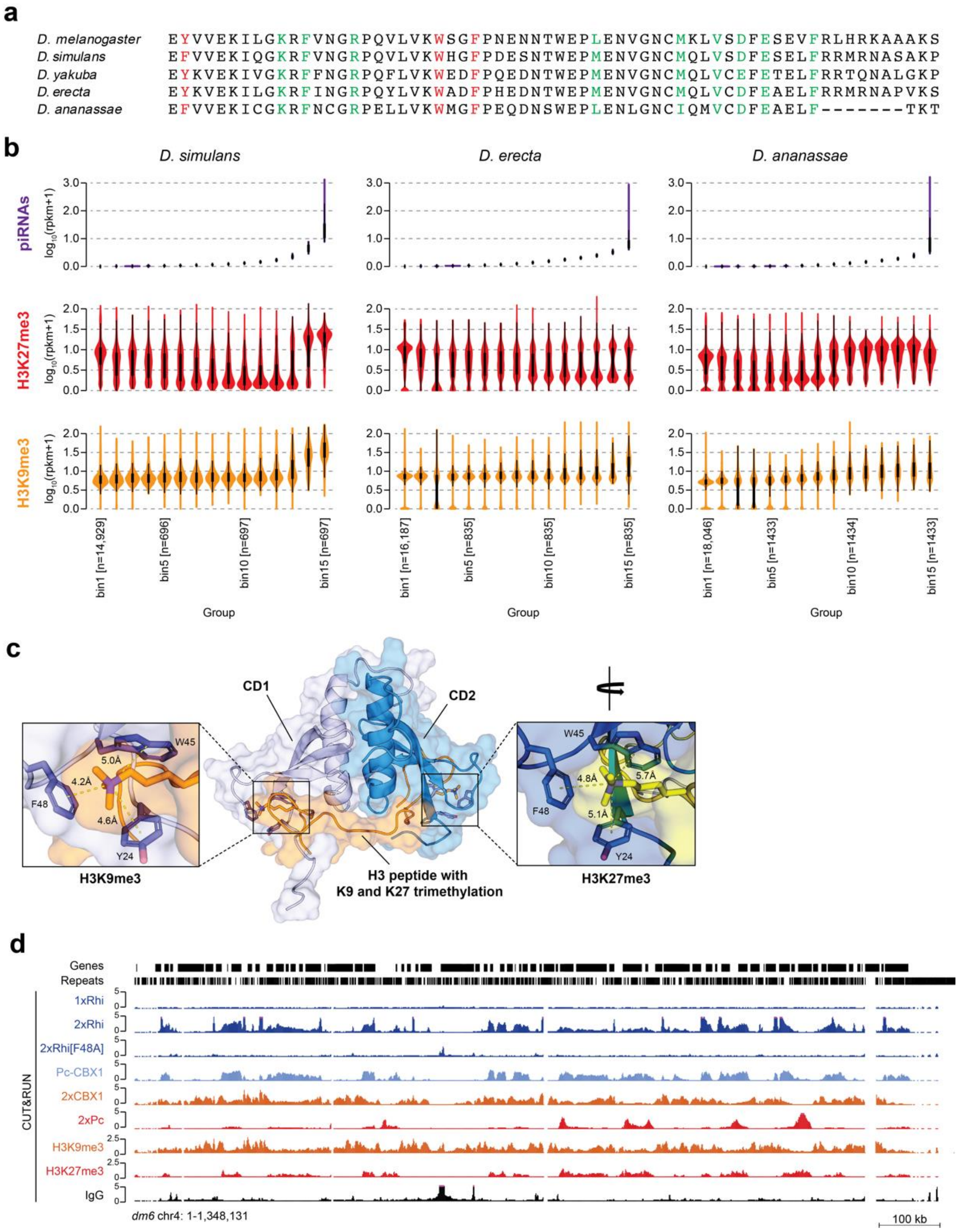


**Extended Data Fig. 8 (related to Fig. 5): Dual-strand piRNA producing loci are associated with both H3K9me3 and H3K27me3 in Drosophilids.**

**a,** Protein sequence alignment of the chromodomains of Rhi orthologs across the indicated species. Residues forming the hydrophobic pocket for trimethyl lysine binding are indicated in red. Residues important for chromodomain dimerization are indicated in green. **b**, Violin plots illustrating the distribution of piRNAs, H3K9me3 and H3K27me3 CUT&RUN signal [$\log_{10}$(rpkm+1)] in *D. simulans*, *D. erecta*, and *D. ananassae* ovaries. All 10 kb bins across the genome were divided into 15 groups. Group 1 contains all 10 kb bins with no piRNA expression. The 14 remaining groups contain an equal number of 10 kb bins. Groups are ranked according to piRNA expression levels with the lowest level of the two strands shown to focus on dual-strand regions. **c,** Model of the Rhi chromodomain dimer in complex with a histone 3 peptide with trimethylated K9 and K27 residues. Representative frame obtained from 1 µS molecular dynamic simulations show the potential for the dimer's interaction with both H3K9me3 and H3K27me3 to be simultaneous. Zoom-in indicated aromatic residues that interact with the methylated lysine group. **d**, UCSC genome browser tracks of *D. melanogaster* chromosome 4 displaying the binding profiles of the indicated chromodomain constructs measured by CUT&RUN (counts per million [cpm] across pooled replicates). Gene tracks are shown above.

**Supplementary Note 1 (related to Extended Data Fig. 1): PRC2 components recovered in screen**

Depletion of *caf1-55* resulted in rudimentary ovaries rendering the analyses of the germline TE expression difficult. Knockdowns of *esc*, *esc1* and *Su(z)12* did not result in sterility of the F1 generation contrary to expected; knockdown of *E(z)* resulted in F1 sterility in the mini screen. This suggests that knockdown efficiencies were not optimal and/or a redundant role of the homologues esc and esc1 and resulted in different effects on TE de-repression; different TE families appear to have different sensitivity to low levels of PRC2.

**Supplementary Note 2 (related to Fig. 2): Antibody cross-reactivity**

Early on in our study we noticed that regions with very high H3K27me3 levels also displayed some H3K9me3 signal. This was particularly evident when plotting H3K27me3 against H3K9me3 signal across genome-wide 1 kb bins (**Fig. N1**), where surprisingly, high H3K27me3 in euchromatic regions is reflected by elevated H3K9me3 signal. A similar observation was made genome-wide (**Extended Data Fig. 3a**), where regions with very high H3K27me3 level also show distinctive H3K9me3 peaks. Since the H3K27me3 methyltransferase E(z) is not expected to be able to write both marks, we concluded that the H3K9me3 antibodies likely displayed a weak cross-reactivity with H3K27me3.

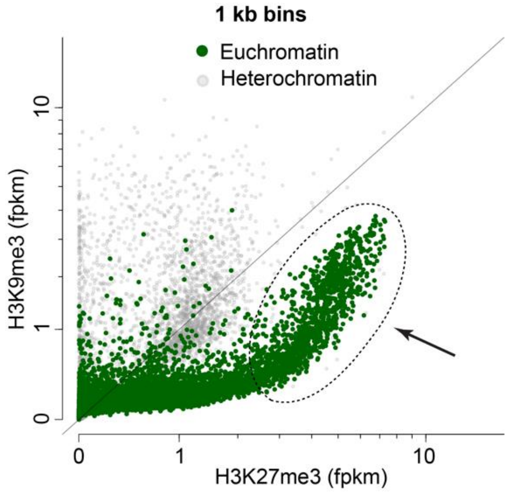


**Fig. N1:** Scatter plot showing H3K27me3 and H3K9me3 levels from CUT&RUN with antibodies in 1 kb bins separated into euchromatin (green) and heterochromatin (grey).

This cross-reactivity was further supported in S2 cells using an *E(z)* knockdown. Globally, S2 cells displayed a marked reduction in H3K27me3 signal following *siE(z)* treatment, as expected, whereas H3K9me3 signal was largely unaltered (**Fig. N2**). However, specific regions with extremely high H3K27me3 signal lost their corresponding H3K9me3 peak following E(z) depletion (**Fig. N2**, yellow areas). Notably, these regions were not bound by the CBX1 chromodomain, a known H3K9me3 reader, and are therefore unlikely to reflect true H3K9me3 signal. Hence, we concluded that H3K9me3 antibodies display a weak cross-reactivity with H3K27me3.

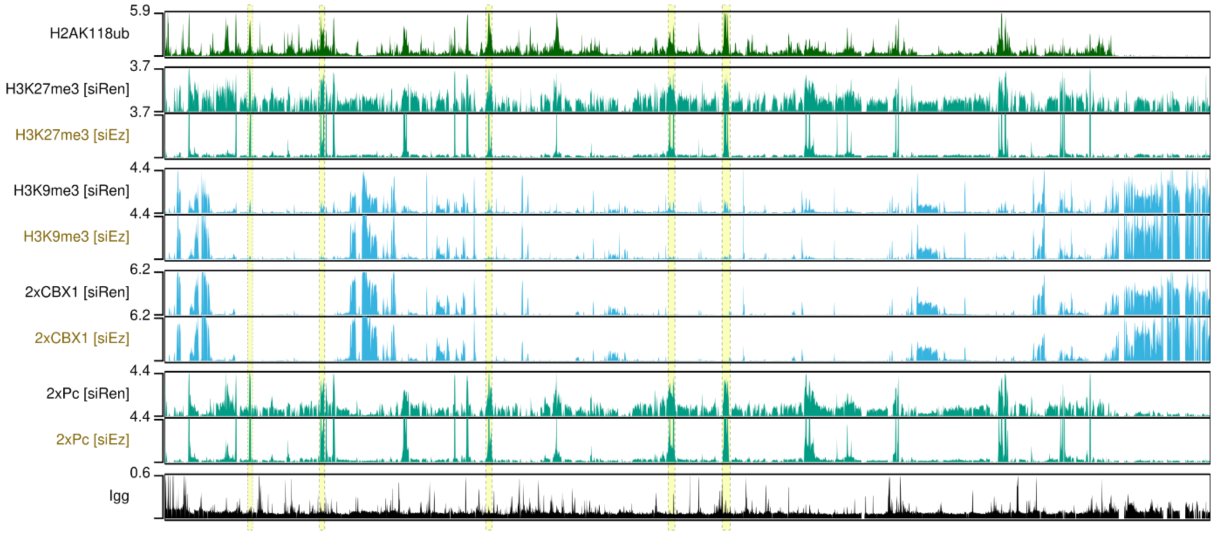


**Fig. N2:** UCSC genome browser tracks displaying H2AK118ub, H3K9me3 and H3K27me3 histone mark signal as well as the 2xCBX1 and 2xPc CD construct signal obtained from CUT&RUN upon the indicated knockdowns.

Although the H3K9me3 and H3K27me3 signals detected by the H3K9me3 antibody strongly differed in amplitude, the potential cross-reactivity made it difficult to evaluate how well H3K9me3 on its own was predictive for dual-strand cluster locations. For these analyses, we therefore opted to use HP1a binding as an alternative method to measure H3K9me3 in *Drosophila* ovaries (**Fig. 2d-f, Extended Data Fig. 3e**), and $CD^{2xCBX1}$ binding as a marker for H3K9me3 in S2 cells (**Extended Data Fig. 7**), ensuring that the results could be unambiguously interpreted.
